# The ecology and adaptive function of clownfish color patterns

**DOI:** 10.1101/2025.08.27.672698

**Authors:** Catheline Y.M. Froehlich, Eleanor M. Caves, Jolyon Troscianko, Miranda Gibson, Tommaso Chiodo, Aurelien De Jode, Korrie Brown, Nina Luckas, Benjamin M. Titus

## Abstract

Anti-predator coloration strategies vary widely from conspicuous to cryptic, and everything between. Coloration serves multiple functions, and determining the selective pressures that drive this diversity is a long-standing mystery. Clownfishes display enormous inter- and intra-specific variation in coloration without the influence of sexual selection, providing an excellent system in which to explore these drivers in closely related and sympatric species. Across clownfishes, three color-pattern phenotypes have evolved convergently with host anemone use, implying adaptive functions linked to the host’s physical and chemical properties. Here we conducted a comprehensive study to infer the ultimate functions of clownfish color patterns. By integrating multiple levels of biological organization, including behavior, visual ecology, dietary niche, and microbiome, our data overwhelmingly indicate that specialist and generalist clownfish color patterns have different underlying ecologies and adaptive functions. Specialist clownfishes background match their anemones, remain near hosts, and have specialized diets and microbiomes. Generalists are highly contrasting with their anemones, remain far from hosts, and have variable diets and microbiomes. Taken together, specialist color patterns align with functional expectations of camouflage while generalists align with aposematic or disruptive coloration. Our study provides novel insights into how ecological conditions shape the evolution of multiple anti-predator coloration strategies.

## Introduction

Among the most remarkable phenotypic traits in the animal kingdom are color patterns used to minimize predation. Known collectively as protective coloration, these traits are used alongside body shapes and behaviors to collectively serve functions ranging from crypsis to mimicry and conspicuous warnings^1^. Yet discerning the ultimate function of an animal’s coloration^1,2^ and the underlying processes that lead closely related species to exhibit different protective strategies remain major challenges^3,4^. This difficulty is compounded by coloration often serving multiple functions simultaneously and therefore comes under diverse selective pressures^2^. Animals considered brightly colored but not sexually dimorphic nor obviously toxic are particularly puzzling^5^, as they do not conform to classic color pattern frameworks^6,7^. In such cases, considering an animal’s ecology is critical to effectively disentangle the adaptive function of coloration, and doing so across species can uncover the underlying processes that generate phenotypic diversity. This approach is valuable because different coloration strategies will functionally be associated with different ecologies, and therefore have testable, ecology-based hypotheses^1^.

One of the most recognizable color patterns on the planet belongs to clownfishes, yet the ultimate function of their coloration remains unresolved. Clownfishes are sequential hermaphrodites that are restricted to a single host anemone for life, and thus do not exhibit sexual dimorphism for color nor sexual selection^8,9^. Multiple hypotheses have been proposed for the function of their color patterns, i.e. inter and intra-specific communication^10,11^ and aposematic warnings of anemone toxicity^12^, without consideration for their phylogeny.

However, a recent study revealed that all clownfishes (28 extant species) have converged upon three distinct color patterns based on host anemone use: orange with one bar (*Radianthus magnifica* specialists), red with one bar (*Entacmaea quadricolor* specialists), and black with multiple thick bars (host generalists)^5^. Thus, we hypothesize that the ultimate function of clownfish color patterns is protective, but the form of protection differs based on their host use. To properly investigate this hypothesis, multiple ecological processes related to host use must be studied in tandem^2^ because the form of protection provided by each color pattern will affect how far clownfish stray from their host and whether predators can see the clownfish in their host, which will impact their prey access and the degree to which host and clownfish microbiomes converge^13^ (Fig 1).

**Figure 1 and Graphical Abstract.**
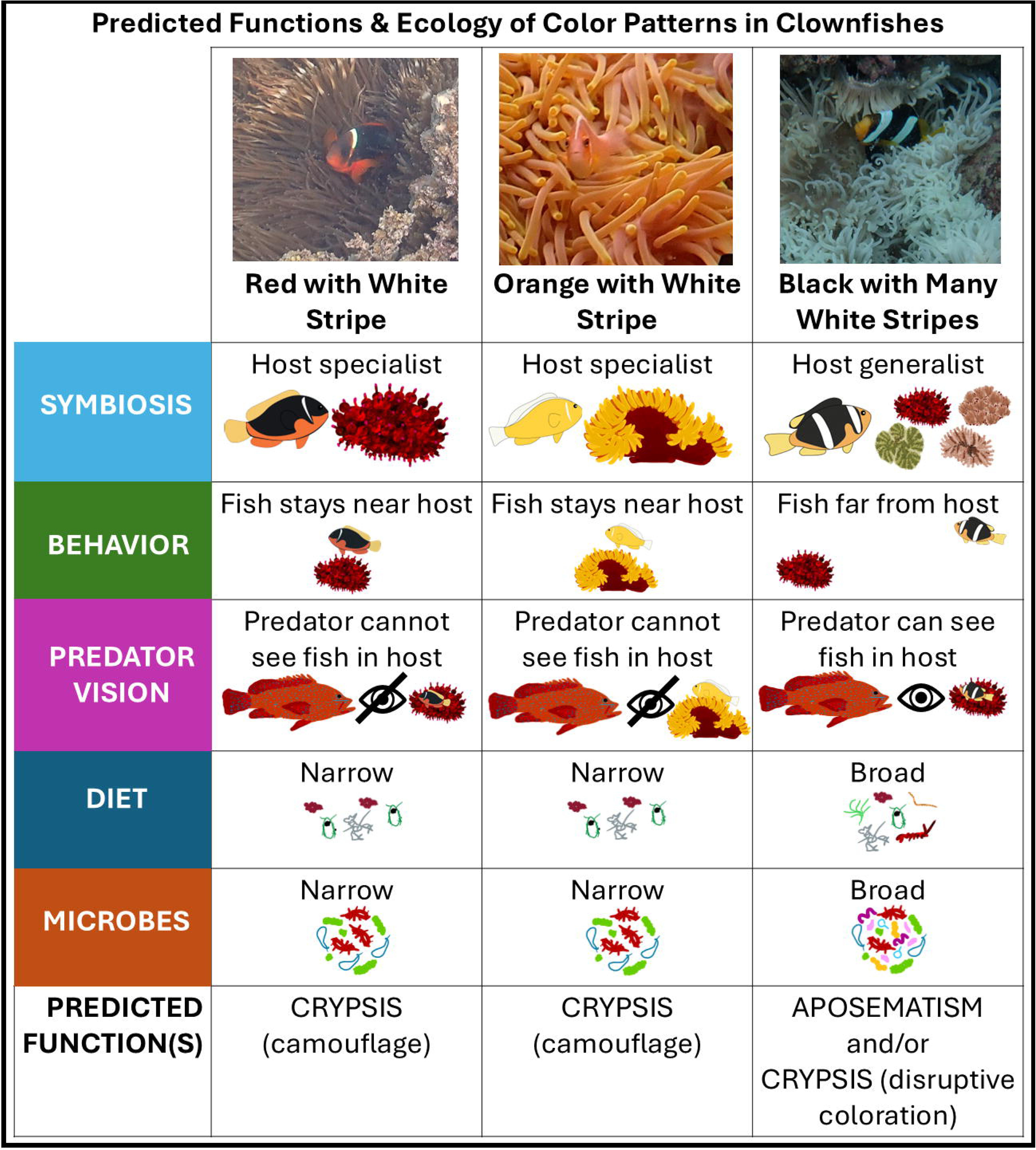
Predictions of the ecology of clownfish color patterns. *A priori* predictions about the ecological context of the three distinct color patterns in *Amphiprion* clownfishes.

Accordingly, we conducted a highly integrative study of three co-occurring clownfish species in Western Australia, each exhibiting one of the three color patterns (Fig 1), two being host specialists and one a host generalist. We predicted the color patterns of both host specialists to serve as camouflage; thus both species would exhibit similar ecologies due to the shared selective pressures of specializing in and around a single host species (Fig 1). Conversely, we predicted that the highly striped color patterns of generalists would serve as disruptive coloration (heavily striped bodies in motion confuse predators)^14–16^ or aposematism (black melanistic coloration signals toxin)^2^ to allow for an ecology that is less host attached. Our findings provide the first evidence that clownfish color patterns generate different protective coloration strategies within closely related taxa.

## Results

### Clownfish behavior and host use

From underwater videos, we quantified the amount of time *Amphiprion clarkii* (host generalist), *A. perideraion* (*Radianthus magnifica* specialist) and *A. rubrocinctus* (*Entacmaea quadricolor* specialist) spent in contact with their host anemones and how far fish swam from their host. We measured distance to the anemone in fish body depths as it is a biologically relevant measure that takes into account fish body-size and life stage. We analyzed the movement behavior of N = 148 fish (21 *A. clarkii*, 18 *A. perideraion*, 109 *A. rubrocinctus*) across species and life stages. The generalist *A. clarkii* moved significantly further from their host anemone than both specialists (p < 0.001; see supplementary material for all statistical analyses), and there were significant differences among clownfish life stages, both within and between species (p < 0.001, Fig 2). Adult *A. clarkii* were rarely in their host anemone and spent over half their time at least twenty body depths away. Conversely, adult *A. rubrocinctus* and *A. perideraion* were either in their host or within one body depth >75% or >90% of the time, respectively. There was very little difference in patterns of host use across life stage in host specialist clownfish as all specialists stay near their host at all times (Fig 2). Subadult and juvenile *A. clarkii* in contrast spent more time in, or close to their hosts than adult *A. clarkii*. Yet subadult *A. clarkii* spent more time out of, and further from, their anemone hosts than either adult host-specialist clownfish.

**Figure 2.**
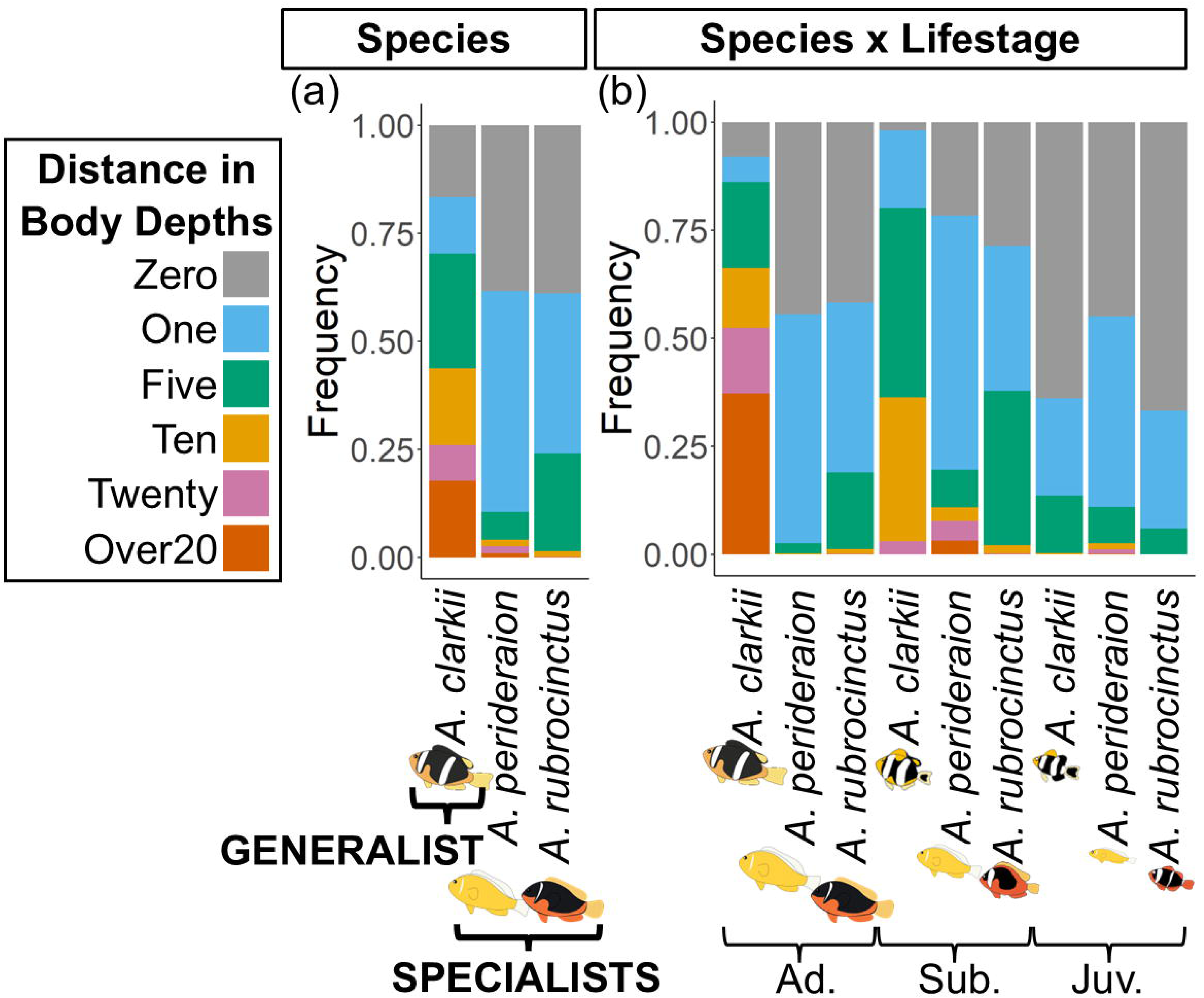
Movement behavior of clownfishes. Frequency of movement behavior of *Amphiprion* species based on species and life stage (N = 148 fish). Movement behavior was measured in distance in body depths away from the anemone. Note: Spp. = species, Ad. = adult, Sub. = subadult, Juv. = juvenile.

### Predator vision and clownfish color-pattern detectability

From the perspective of a human visual system, the generalist *A. clarkii* appear to have multiple thick white stripes on a largely black background throughout ontogeny. *Amphiprion perideraion* maintain a single stripe on an orange body throughout ontogeny, while *A. rubrocinctus* transitions from three stripes as juveniles to a single stripe as adults on a red body. However, to understand the adaptive function of clownfish color patterns, we must account for the visual system of the intended receiver in nature, which can differ significantly from that of humans. Therefore, using color-calibrated underwater photography for host sea anemones and their clownfishes, we modelled the conspicuousness of clownfish color patterns against their host background from a predator visual system. We used the visual physiology of the coral trout *Plectropomus leopardus*, a common piscivorous reef predator, which has published measures of spectral sensitivity and acuity for coral trout^17^ and existing measures of weber fraction from another species with similar vision (triggerfish)^18^. For N = 53 fishes (14 *A. clarkii*, 14 *A. perideraion*, 25 *A. rubrocinctus*), we modelled aspects of both luminance (i.e. brightness, using stimulation of the medium-wavelength cone) and color vision at three different distances to mimic predator striking (0.4m and 1m) and searching (5m) distances, specifically quantifying how different clownfish color patterns were from their anemone backgrounds. For differences in color pattern (quantified in units of ‘just noticeable difference’), we considered values at or below 1 to indicate that a clownfish was indistinguishable from its anemone at that distance.

At all distances and life stages, the host generalist *A. clarkii* was more luminant against their host anemone backgrounds than either host specialist clownfish *A. perideraion* or *A. rubrocinctus* from the predator perspective (i.e. they were brighter and more detectable by the predator *P. leopardus,* Fig 3a, p < 0.001). At both striking distances and all life stages, the color patterns of generalist *A. clarkii* were more distinguishable from their anemones than both host specialists from the predator perspective (Fig 3b, p < 0.001). From a searching distance, there were minimal differences between the color patterns of generalists and specialists compared to their anemones, but distinct difference were still observed in their luminance (Fig 3b).

**Figure 3.**
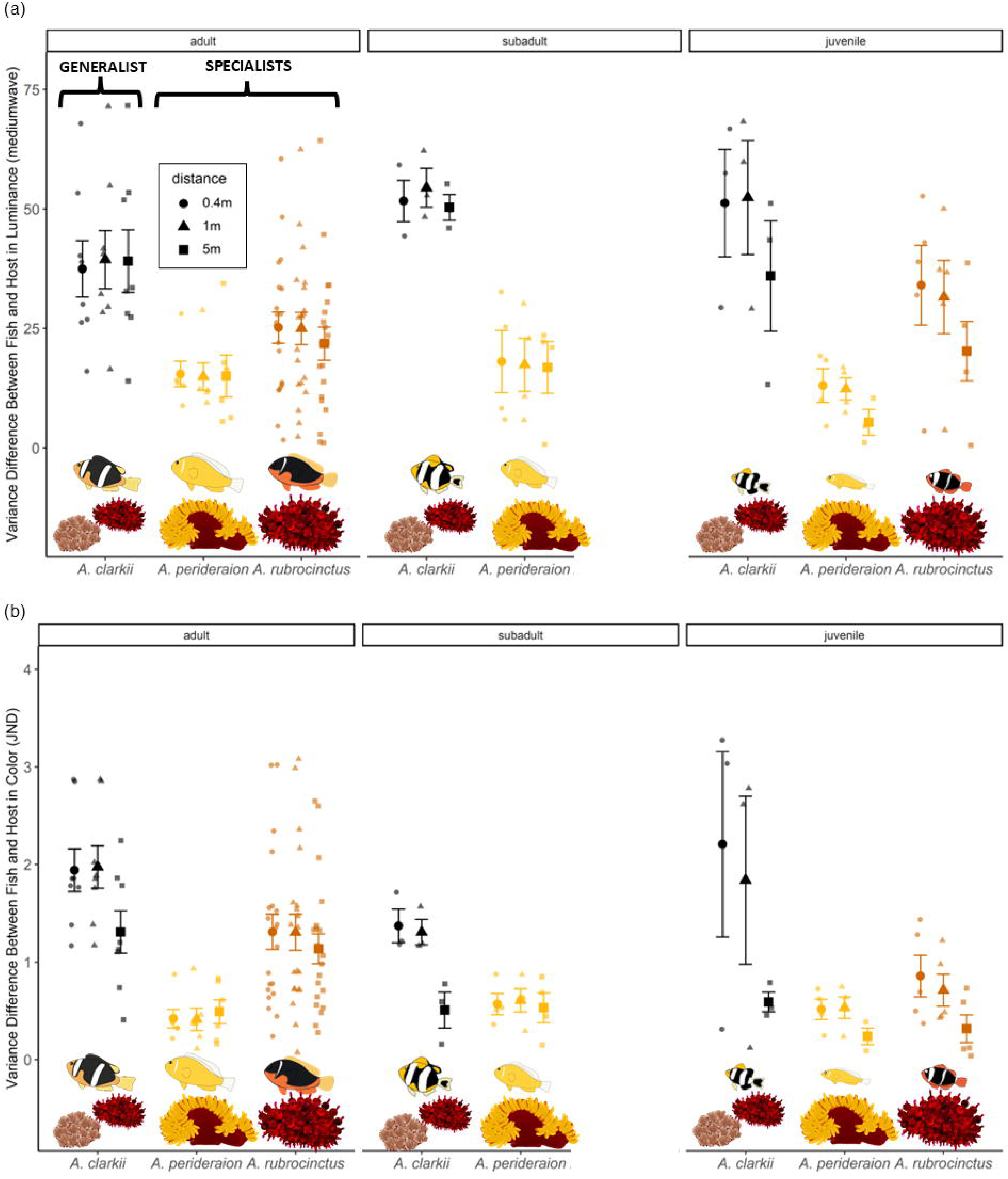
Visual conspicuousness of clownfish color patterns. Predator differentiating between clownfish and host anemones via (a) luminance pattern energy (i.e. variance) at different distances, (b) color pattern energy (i.e. variance of just noticeable difference) at different distances (N = 53 fish, N = 30 anemones). JND = just noticeable difference. The dots with error bars represent mean and standard error, respectively.

### Dietary niche

We collected and analyzed δ^13^C and ^15^N stable isotope signatures for N = 45 clownfishes (14 *A. clarkii*, 6 *A. perideraion*, 25 *A. rubrocinctus*) in two adjacent regions within the study location. The dietary niche of generalist *A. clarkii* was much larger than both specialists (p < 0.01), which were similar in size (Fig 4a). The majority of the variation was with δ^13^C signatures, as *A. clarkii* ate a more pelagic or benthic diet depending on region, indicating a generalist diet, whereas there was no difference in region for the specialist (Suppl Fig 21). Generalist *A. clarkii* and specialist *A. perideraion* had slightly higher δ^15^N signatures than specialist *A. rubrocinctus* (Fig 4a; trophic position mean ranged from 2.6 to 3.1; p < 0.01), although there were no regional differences in δ^15^N signatures (Suppl Fig 21). Among clownfish life stages, there was no overlap in dietary niche of the different life stages for either *A. clarkii* nor *A. perideraion*. However, the dietary niche of juvenile *A. rubrocinctus* was almost completely enclosed in their adult niche (Fig. 4b; p < 0.01). Juvenile *A. clarkii* had dietary niche that spanned greater δ^13^C values than juvenile or adult *A. perideraion* and *A. rubrocinctus* (Fig 4b).

**Figure 4.**
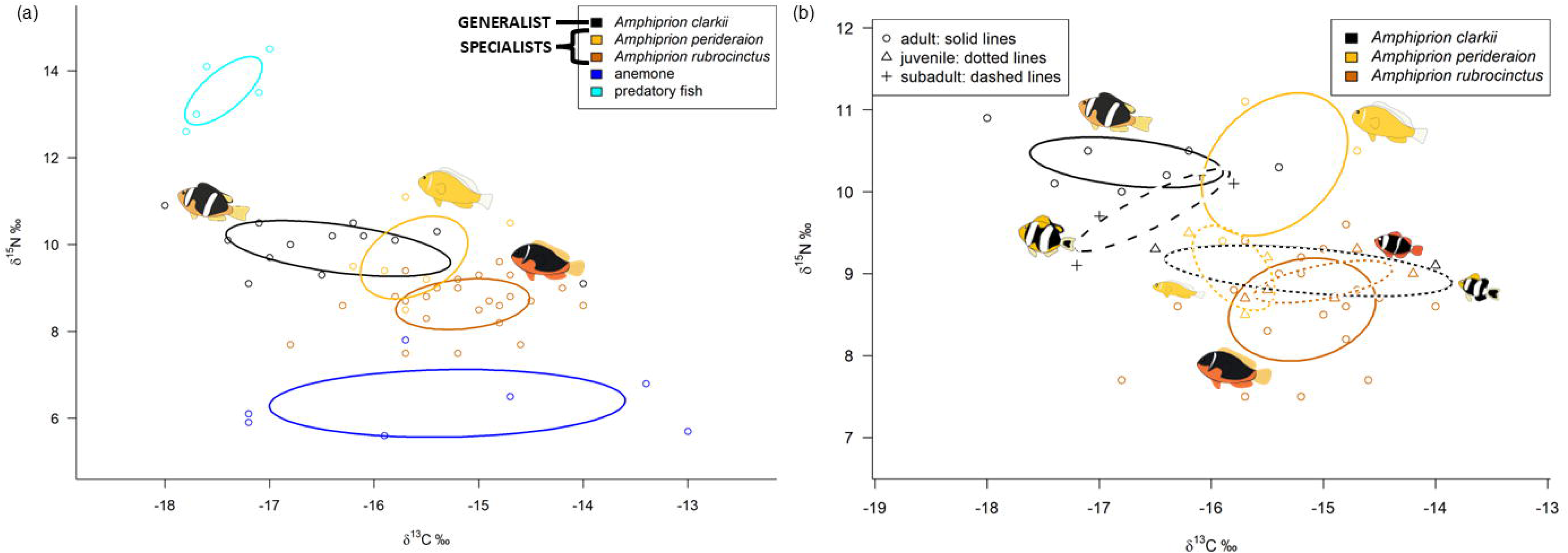
Dietary niche of clownfishes. Isotopic niche overlap for dietary niche of (a) clownfishes (N = 45), pooled anemones (N = 6), and pooled predatory fish (N = 4); and (b) different life stages of clownfish. Standard ellipses areas (40%) are depicted for carbon and nitrogen isotopes.

### Microbiome diversity

We sequenced the microbiome from the mucus and tentacles of anemones and the skin, gills, and guts of the clownfishes. There were 19 anemone mucus samples, 28 anemone tentacle samples, 25 fish skin samples, 30 fish gill samples, and 34 fish gut samples that remained after we called and filtered Amplicon Sequencing Variants (ASVs) with a total of 11,980 microbiome ASVs. There were not enough samples retained of the host anemone *Radianthus crispa* nor different clownfish life stages to include in the analysis. The generalist clownfish *A. clarkii* had greater microbiome community variation than either specialist *A. rubrocinctus* or *A. perideraion* (Fig 5; p = 0.03). Although specializing on different hosts, there was more overlap in the microbiome communities of both specialists than there was between either specialist and the generalist *A. clarkii*. Host anemone microbiomes (*R. magnifica* and *E. quadricolor*) were fully enclosed within, smaller and less variable than any of the clownfish microbiomes (Fig 5).

**Figure 5.**
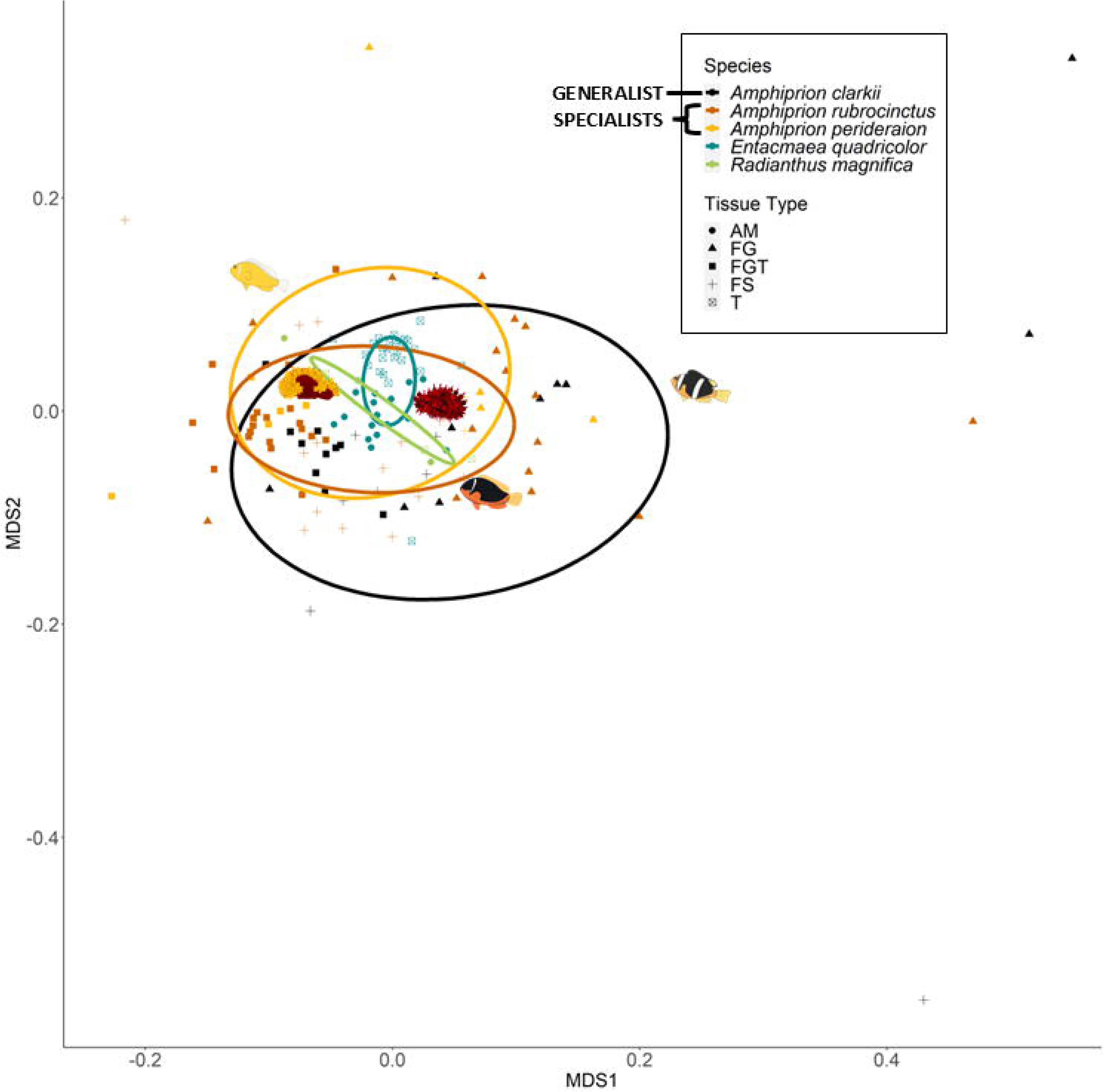
Microbiome of clownfishes and anemones. Non-Metric Multidimensional Scaling (nMDS) plot of the microbiome of species (colors and ellipses) by tissue type (shapes). AM = anemone mucus (N = 19), FG = fish gills (N = 30), FGT = fish guts (N = 34), FS = fish skin (N = 25), T = anemone tentacles (N = 28).

## Discussion

Our findings support the hypothesis that the ultimate functions of clownfish color-patterns are protective and directly linked to the way host specialist (*Amphiprion perideraion* & *A. rubrocinctus*) and generalist (*A. clarkii*) species interact with their host sea anemones. Although specializing on distantly related host sea anemones, the underlying ecology and adaptive function of both specialist color patterns have converged. Specialists remain extremely close to their long-tentacled host anemones at all life stages and have more specialized dietary niches and microbiomes than generalists. Through a loss of thick white vertical bars and reduced melanin production, specialist clownfish color patterns are virtually indistinguishable to potential predators against their host sea anemones at both striking and searching distances. Specialist color patterns therefore function as background-matching camouflage. In contrast, the generalist clownfish *A. clarkii* ventures far from their host sea anemones and have a broader dietary niche and microbiomes than specialists. They have highly contrasting color patterns that are easily detected by predators against their anemones. Our data supports the hypothesis that *A. clarkii* color patterns function as either aposematic (black melanistic coloration signals toxin) or disruptive coloration (heavily striped bodies in motion confuse predators). This enables generalist clownfishes to explore and forage in diverse environments while deterring or confusing predators. The integrative ecological approach we use here reveals how species from a co-occurring taxon with seemingly overlapping microhabitat requirements have converged upon three distinct color patterns that each serve a different form of protective coloration. Our data provide the first unifying explanation for the ultimate adaptive functions of clownfish color patterns.

Clownfish color patterns are often described as “brilliant” and “highly visible.” Our findings suggest that only color patterns from host generalist species can be described in this manner^5^ when the relevant viewer (e.g. predator) is considered with the relevant natural background (i.e. the anemone). Phylogenetic reconstruction has shown that common ancestor of all clownfishes was likely a generalist with black base body color and possessing 3 white vertical bars^10,11,19^. Based on our ecological data, ancestral generalist clownfishes, just like extant species, probably spent little time within their hosts sea anemones past the juvenile life stage, foraged broadly as indicated by their broad dietary niche, and encountered diverse coral reef microhabitats that led to highly variable microbiomes. Their behavior, combined with the ability to live within any host sea anemone regardless of tentacle density, morphology, and toxicity, resulted in the evolution of a protective phenotype that was effective in any host. As previously suggested for clownfish^12^, monarch butterflies and venomous snakes^20–22^, their striped and melanistic bodies may signal aposematism since their host anemones are toxic^23^. Generalist clownfish may therefore be incorporating the host toxin into their tissues, as seen in caterpillars^4^ that live on toxic hosts or simply signalling their host’s toxicity^12^. However, adult and subadult generalists rarely spend any time interacting with their host, which reduces the potential for the anemone toxin to be incorporated into the fish. Alternatively, their stripes may confuse predators on their actual location via disruptive coloration. By using specific movements over different backgrounds, the stripes in motion can make the fish cryptic, as seen in some lizards^24^, snakes^25^, zebras^14^, bats^26^ and humbug damselfish^16^. Our data are consistent with the hypothesis that generalist phenotypes are either aposematic or a form of disruptive coloration, or both, and further research will be important for differentiating between the two.

In contrast, the ecology of both host specialists converged similarly and suggests a single antipredator strategy to conform to the host anemone. Both specialists are not conspicuous to predators against their host anemones, stay close to their hosts, and exhibit more specialized dietary niches and microbiomes than generalists. Some specialist clownfishes have the ability to swim far from their host as per swimming performance and muscular morphology, however they choose to stay close to their host^27^. Such an ecology suggests a strongly site attached relationship with their host in order to camouflage within their long tentacles, as seen in nontoxic caterpillars^4^, Chrysomelinae beetles as determined with their specialized diet^28^, and eastern fox squirrels^29^. The only marked difference is that juvenile *A. rubrocinctus* were more easily distinguished against their hosts than juvenile *A. perideraion*, as they have multiple stripes that are lost with ontogeny^10,11,30^. The heavily striped bodies of *A. rubrocinctus* juveniles may either serve for rank recognition^31^ or disruptive coloration as secondary functions, or may be a remnant of their common ancestors^5^, who were host generalists. Specialists do not exhibit aposematic signals as they are hard to differentiate from their hosts, and stay near and within their hosts, which have the longest and most densely packed tentacles to allow for full concealment. Thus, the ecology of both specialists confirms that their color patterns have converged as a means of camouflage to background match and blend into their hosts.

Our work underlines the need to assess multiple ecological processes in tandem to properly investigate the adaptive function of animal color patterns, particularly in co-occurring taxa. Several studies have used ecological context to provide new hypotheses for color pattern function in songbirds, lizards and beetles^2,3,28^. As seen in other taxa^20,21^, no single function explains the diversity of clownfish coloration. Much clownfish literature has focused on the hypothesis that gaining or losing stripes throughout ontogeny of some clownfish species is a means of intraspecific signing for social rank recognition and/or interspecific signing for species recognition^10,11,30^. Additionally, studies have found that clownfish can distinguish social ranks using ultraviolet reflectance^32,33^, though clownfish predators (like coral trout *Plectropomus leopardus* used in our vision model) rarely possess ultraviolet sensitivity^34^. However, intraspecific and interspecific communication cannot account for color pattern evolution across the clade nor color pattern convergence with the host anemones^5^, and instead are likely secondary functions of clownfish color patterns. Thus, our study is the first to disentangle the ultimate function of clownfish color patterns and show that different forms of antipredator coloration have evolved based on host anemone use within a single clade.

## Online Methods

### Sample Sites and Collection at Research Station

The study was completed in Coral Bay, Ningaloo Reef, Australia (-23.140419, 113.769153) in June 2023. Clownfish were haphazardly encountered among reefs north and south of the Coral Bay Research Station. There was an anoxic event in 2022, which resulted in coral death in both regions but the southern region had less live coral cover in 2023. *Amphiprion perideraion* colonies were extremely rare (n = 3) and had at least 4 individuals that were found on a single *Heteractis magnifica* anemone. *Amphiprion rubrocinctus* (n = 21 colonies) were either alone or in colonies with up to twelve individuals and were only found on *Entacmaea quadricolor* with multiple anemones clustered together. *Amphiprion clarkii* (n = 13 colonies) were either alone or in colonies up to six individuals and were found in both *E. quadricolor* and *Heteractis crispa*. On several occasions (n = 6), *A. clarkii* and *A. rubrocinctus* were cohabiting in mixed species colonies on *E. quadricolor.* When *H. magnifica* or *H. crispa* were found, they were usually on top of coral rubble or in sand, respectively. When *E. quadricolor* were found, they were clustered with many clonal individuals embedded within coral rubble with only tentacles sticking out.

For each clownfish colony, cameras were deployed on tripods to observe movement behavior. For videotaping, GoPro 7 cameras were placed with extended battery packs in custom waterproof housings that were built at Dauphin Island Sea Lab for long-term footage. Depending on the size of the colony, either one or two cameras were placed in order to videotape the whole colony. After videotaping, water samples were collected with 5L collapsable water bottles. Tentacle clippings were collected with tweezers and anemone mucus was collected using 60ml Loctite syringes. One adult and one juvenile (if present) were collected from each colony and species, otherwise a subadult was collected if adults were not present. Fish were collected without anaesthetic using hand nets and placed in Ziploc bags. Underwater, the fish within bags were placed in front of a foamed ethylene-vinyl acetate sheet (common mousepad) and DGK Kustom Balance color checker card. Fish were photographed in relatively even lighting (no flash; no direct sunlight) to limit over exposure^35^ with Olympus TG-6 and Sony A1 cameras set to photograph in RAW format. Several photographs were taken since even lighting and no turbidity were difficult to achieve underwater. Anemones were similarly photographed. All the videotaping, photographing and collections were completed on scuba. Then fish were brought to the surface and their whole body was swabbed with two sterile cotton swabs and stored in RNA later. Fish were then placed in ice buckets for euthanasia.

Samples were brought back to Coral Bay Research Station for processing. Water samples were filtered onto 2 µm filter paper with the eDNA Citizen Science Sampler (Smith-Root). Filter paper was stored in 2ml tubes in a -20°C freezer. Fish were dissected to collect fish gills, guts, and muscles and then the body was stored in 70% ethanol. Anemone mucus was placed into 50ml falcon tubes and the mucus was allowed to settled out of the seawater for 5-10min. The mucus was then pipetted into 2ml tubes. Anemone tentacles (half of each sample), mucus, fish gills and guts were stored in RNAlater for microbiome analysis. Fish muscles and anemone tentacles (other half of sample) were placed in distilled water for 30min, then dried for 24-48 hrs at 70°C in a drying oven, and were stored at room temperature for stable isotopes analysis. Samples from other marine species, including predatory fish, pelagic and benthic invertebrates, and algae were collected *in situ* in order to represent key functional groups (primary producers, primary consumers, secondary consumers, top predators). Predatory fish samples were obtained from local sportfishermen. These other marine samples were placed in distilled water for 30min, dried and stored for stable isotopes analysis. All samples and footage were brought to Dauphin Island Sea Lab (DISL), Dauphin Island, USA for further analysis and frozen samples were stored in a -80°C freezer.

### Movement Behavior: shelter proximity

For each colony, the videos were trimmed to remove the first 5min after divers were observed. Then each minute was analyzed for the next 90min for movement behavior of each individual fish in the colonies. Distance of fish to the anemone was measured by calculating how many body depths (i.e. maximum distance between the ventral and dorsal part of the fish) the fish was away from the anemone. Several studies have used actual distance measured through 3D analysis from using multiple cameras at once and special software; however such a measurement does not consider how a specific distance may represent different effort or risk for a smaller versus larger fish. Although absolute distance could be calculated by dividing the distance by fish size, each fish would need to be collected to measure their size. Instead, the distance measured in body depths is a measurement directly related to the size of each individual fish that can be measured in video footage alone (whether 2D or 3D), minimizing fish handling. Although total body length (tip of snout to end of tail) is the typical measurement of fish size, body length cannot be measured in most frames of video because fish are generally turning and not horizontally flattened. Therefore, distance in body depth is a biologically relevant measurement to investigate how far fish move, and it can be measured in each frame of any multidimensional footage with any video software. To calculate the distance to anemone in body depth at each minute interval (equation (1)), a ruler was each to measure the distance to anemone and body depth of the fish in the frame of each minute interval (*i*).

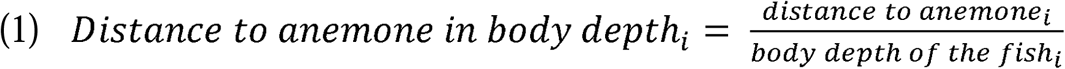

Distance in body depths per interval were binned in categories based on predation risk via distance to shelter (anemone): zero (in shelter; at least 1/4 of fish is inside the anemone), one body depth away (high concern for retreating to shelter), five body depths away (medium concern for shelter), ten body depths away (low concern for shelter), twenty (very low concern for shelter), and over twenty body depths away (little to no concern for shelter). When fish were out of the anemone, we also noted whether the fish was either above the anemone or in the periphery/under the anemone. Fish were removed from analysis if their location could not be identified for at least half of the time (this removed 8 individuals from analysis). For data analysis, the distance in body depth was analyzed using a cumulative linked mixed model with the following factors: species (fixed factor), life stage (fixed factor), region (north or south; fixed factor), type (single or mixed species colony; fixed factor), fish number (random factor), and colony (random factor). The analysis was completed in R v4.2.2^36^ with the ordinal^37^, rcompanion^38^, and tidyverse^39^ packages.

### Antipredator Coloration: predator vision

Image analysis on RAW photographs was completed in micaToolbox v2.3^35,40^ to determine whether fish could be discriminated from their anemone by a common predator, the coral trout *Plectropomus leopardus*. We used two color checkers for our study: an Xrite ColorChecker classic card (not waterproof) to calibrate the cameras used underwater, and an underwater DGK color checker to take photographs of fish and anemones underwater. To measure the reflectance of each square in our color checkers, we used an Ocean Optics USB-2000 Fiber Optic Spectrophotometer, which was calibrated with a Spectralon 99% white standard. To calibrate the camera for image analysis within micaToolbox, the Xrite color checker was photographed on a plain gray background in cloudy conditions with each camera used underwater. To select the best underwater photograph of each animal, all photographs were screened to check for overexposure, turbidity, and visibility of gray scale boxes on the DGK color checker. The best photograph was loaded as a linear normalized multispectral image and converted into a cone catch image with the coral trout D65 vision model. For predator vision analysis, the 0.5 weber fractions of a generalist teleost fish (triggerfish 1:2:2)^18^ were selected since no weber fraction data exists for coral trout. However acuity has been previously measured in the coral trout at approximately 10 cycles per degree^17^, so that value was used to represent predator acuity. Accordingly, Quantitative Colour Pattern Analysis (QCPA) analysis was completed for coral trout vision at 5m, 1m, and 0.4m to mimic predator vision at different distances. The mean and standard deviation of luminance for the medium wavelength-sensitivity cone of the animal color pattern was measured. The difference in these values between the clownfish and anemone was calculated using equation 7 from^41^ for mean (average luminance) and standard deviation (luminance pattern energy) values separately. The receptor noise limited (RNL) model was used to calculate Delta-S, a measure of chromaticity distance, and from this we calculated the just noticeable difference (JND) between clownfishes and anemones by taking the Euclidean distance for mean (color, average JND) and standard deviation (JND pattern energy) values separately.

### Diet Ecology: stable isotopes

After samples were transported to DISL, they were dried again in a drying oven for up to 24hrs at 70°C to remove any humidity from transportation. All samples were powdered either by hand with a metal spatula if there was very little tissue (< 10mg), or using a Bead Ruptor 12 (Omni International) if there was enough tissue (≥ 10mg). For each invertebrate sample, half of the sample was lipid treated with a 2:1 chloroform:methanol solution by modifying previous protocols^42,43^: add solution to cover tissue completely, vortex for 1 min, add a few drops of distilled water, vortex for 1 min, centrifuge for 10 min at 10,000 rpm, decant supernatant, repeat all steps, then let evaporate overnight in fume hood at room temperature. For each sea star sample, half of the sample was acid treated due to their calcareous characteristics by modifying a protocol^44^: add 1 ml 1N HCl, vortex, let sit for 5 min, add 1 ml 1N HCl, vortex and let sit until no more effervescence or repeat as needed, centrifuge for 5 min at 1800 rcf, decant supernatant, repeat all steps but all solution to sit for 1hr before centrifuging, rinse with 9ml of distilled water, vortex, centrifuge again, and decant supernatant. Each sample was dried again in the drying oven at 70°C and pulverized into a powder if needed. Between 0.314 to 1.586 mg of each sample (both treated and untreated) was weighed in tin capsules (3.3 x 5 mm) using a microbalance and organized in 96-well plates. The plates were sent to The Center for Stable Isotope Biogeochemistry (CSIB) on the University of California, Berkeley campus for δ^13^C and δ^15^N stable isotope analysis. Isotopic values were denoted in parts per thousand (‰) using the standard δ notation, where δX = [R_SAMPLE_/R_STANDARD_ – 1) × 1000]; with R representing the high to low isotope ratios. Nitrogen values were retained from untreated samples only since lipid and acid treatments may alter nitrogen isotope signatures^45^.

Out of all invertebrate samples that were also lipid treated, only oyster samples had carbon to nitrogen ratios above 4. Therefore, for carbon analysis the lipid treated samples of oysters were used, but non-lipid treated samples were used for other invertebrates. For sea stars, acid treated carbon samples were used. For nitrogen analysis, all untreated samples were used. For stable isotopes analysis, only 7 anemone samples were collected from different species hosting clownfish, thus anemone samples from different species were pooled together since they were only used as reference. Predatory fish samples were also pooled together as there was only one sample per species collected and they were only used for reference.

Isotopic niche space was calculated in the SIBER package^46^ in R v4.2.2^36^ using standard ellipse (40% confidence intervals) area corrected for small sample size (SEA_c_) and fitted with Bayesian models (10^4^ iterations). If there was less than a 5% overlap in SEA_c_ among any groups, the isotopic niche space among those groups was considered significant. If there was more than 60% overlap in SEA_c_, the groups had a shared isotopic niche space^47^. Trophic position was calculated using the tRophicPosition^48^. We used trophic fractionation values that were established for an *Amphiprion* species^49^. A oneBaseline approach with 20^4^ model iterations was used with algal sources collected *in situ* as baseline representations of primary producers (trophic position = 1). Trophic position and δ^13^C values were analyzed using linear models and Tukey’s post-hoc tests for pairwise comparisons for each analysis: (1) piscivorous fish vs. each of the three clownfish species vs. anemones (fixed factor); (2) each of the three clownfish species (fixed factor) by adult and juvenile (a.k.a. life stage; fixed factor); and (3) *A. clarkii* and *A. rubrocinctus* (fixed factor) by region (north or south; fixed factor). Note that subadults were only collected from *A. clarkii* since they were solitary on anemone hosts, therefore they were not included in the linear model for (2). Data was transformed to meet normality and homoscedasticity using Q-Q plot visualization, histograms, and residuals over fitted plots. Outliers were removed if their standard residuals fell outside of 2.5 standard deviations from 0. Additional R packages included: tidyverse^39^, GGally^50^, reshape2^51^, plyr^52^, lme4^53^, emmeans^54^, piecewiseSEM^55^, LMERConvenienceFunctions^56^.

### Microbiome

Once samples were transported to DISL, they were stored in the -80°C freezer before use. DNA was extracted from each sample using the Qiagen blood and tissue kit. The water and mucus samples used twice the amount of mastermix than for other samples in order to get enough extraction for analysis. Several samples of each tissue type were tested via PCR using the 515F-806R primer pair^57–59^ for the 16S region to check that the protocol meets requirements for microbiome analysis. Samples were sent to the University of Maryland Microbiome Service Laboratory to undergo 16S library preparation and Illumina sequencing.

Data underwent demultiplexing, trimming, filtering, denoising, and removal of chimeric samples. Contaminated Amplicon Sequence Variants (ASVs) were identified and removed using the decontam package. Eukaryote ASVs were removed. Via prevalence filtering, ASVs were removed if at least 80% of their abundance came from water samples and less than 20% came from other samples. Samples were removed with less than 3000 sequences and outliers were removed. Using a permutational analysis of variance (PERMANOVA), fish microbiome samples were compared among species (fixed factor) and life stage (fixed factor), without anemone samples. There were not enough samples per life stage to include tissue type in the latter analysis. A separate PERMANOVA was run to compare the microbiome of all fish and anemone samples by species (fixed factor) and tissue types (fixed factor), without life stage as a factor. Following the PERMANOVA, pairwise testing and non-Metric Multidimensional Scaling (nMDS) plots were used to compare and visualize results. The following packages were used in R for the analysis: decontam^60^, pairwiseAdonis^61^, phyloseq^62^, tidyverse^39^, and vegan^63^.

## Acknowledgements

We would like to acknowledge the Baiyungu, Thalanyji and Yinigurdira people as the Traditional Owners of Ningaloo Marine Park. Our project was funding by the National Science Foundation PurSUiT grant to B.M.T. and Estefania Rodriguez (DEB-1934274), University of Alabama start-up funds to B.M.T., and National Science Foundation Postdoctoral Research Fellowship in Biology to C.Y.M.F. (PRFB-2305953).

## Data Availability

All genomic data is available on NCBI project: PRJNA1309112. All other data is available at https://knb.ecoinformatics.org/view/doi:10.5063/F1W957PC.

