## Supplementary Data for "The ecology and adaptive function of clownfish color patterns"

### <u>**Supplementary Data**</u>
### <u>**For Manuscript:**</u>

The Ecology and Adaptive Function of Clownfish Color Patterns

#### <u>Results:</u>

#### ***CLOWNFISH BEHAVIOR AND HOST USE***

Supplementary Table 1. Output of all behavioral stats. CLMM = cumulative linked mixed model.

| Response Variable | Model | Predictor variable | Factor Type | df | Model Statistic | Test-value | p-value |
|---|---|---|---|---|---|---|---|
| Distance in Body Depth to | CLMM | Species | fixed | 2 | $\chi 2$ (chi-squared) | 48.37 | < 0.0001* |
| Anemone | | Fish Number | random | | Sample size | n = 12179 | |
| categorical variable (6 levels) | | Group | random | | | | |
| Distance in Body Depth to | CLMM | Species | fixed | 2 | $\chi 2$ (chi-squared) | 67.63 | < 0.0001* |
| Anemone | | Life stage | fixed | 2 | | 35.05 | < 0.0001* |
| categorical variable (6 levels) | | Species x Life stage | fixed | 7 | | 630.85 | < 0.0001* |
| | | Fish Number | random | | Sample size | n = 12179 | |
| | | Group | random | | | | |
| Distance in Body Depth to | CLMM | Species | fixed | 1 | $\chi 2$ (chi-squared) | 81.66 | < 0.0001* |
| Anemone | | Region | fixed | 1 | | 0.10 | 0.76 |
| categorical variable (6 levels) | | Species x Region | fixed | 1 | | 6.63 | 0.01* |
| Adults Only: *A. clarkii,* and | | Fish Number | random | | Sample size | n = 6659 | |
| *A. rubrocinctus* | | Group | random | | | | |
| Distance in Body Depth to | CLMM | Species | fixed | 1 | $\chi 2$ (chi-squared) | 81.62 | < 0.0001* |
| Anemone | | Type (Single/Multi Species) | fixed | 1 | | 0.01 | 0.92 |
| categorical variable (6 levels) | | Species x Type | fixed | 1 | | 0.15 | 0.70 |
| Adults Only: *A. clarkii,* and | | Fish Number | random | | Sample size | n = 6659 | |
| *A. rubrocinctus* | | Group | random | | | | |

For regional and colony composition comparisons, only adult *A. clarkii* and *A. rubrocinctus* were included due to limited availability. In the north, adult *A. clarkii*, were in the anemone 15% of the time whereas those in the south were less than 5% in the anemone ($p < 0.01$). For adult *A. rubrocinctus*, there were no marked differences. When comparing single species and mixed species colonies (*A. clarkii* and *A. rubrocinctus* cohabitating), fish did not change their movement behavior ($p > 0.38$), although it was much rarer to see *A. clarkii* in their anemones when in mixed species colonies.

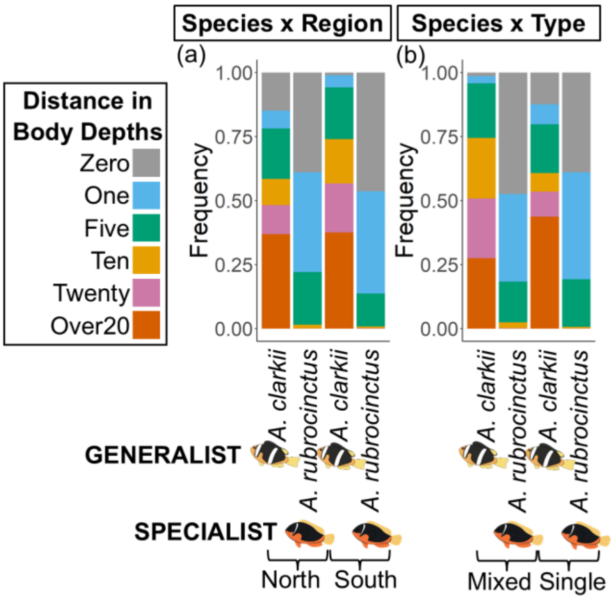


Supplementary Figure 1. Frequency of movement behavior of *Amphiprion* species based on region and type (single species or mixed species colonies). Movement behavior was measured in distance in body depths away from the anemone. Note: Spp. = species, Ad. = adult, Sub. = subadult, Juv. = juvenile. Generalist species = *A. clarkii*, specialist species = *A. rubrocinctus* & *A. perideraion.*

When fish were not in the anemone, they were either above the anemone, as seen all species, or in the periphery of the anemone, as seen consistently only in *A. perideraion*.

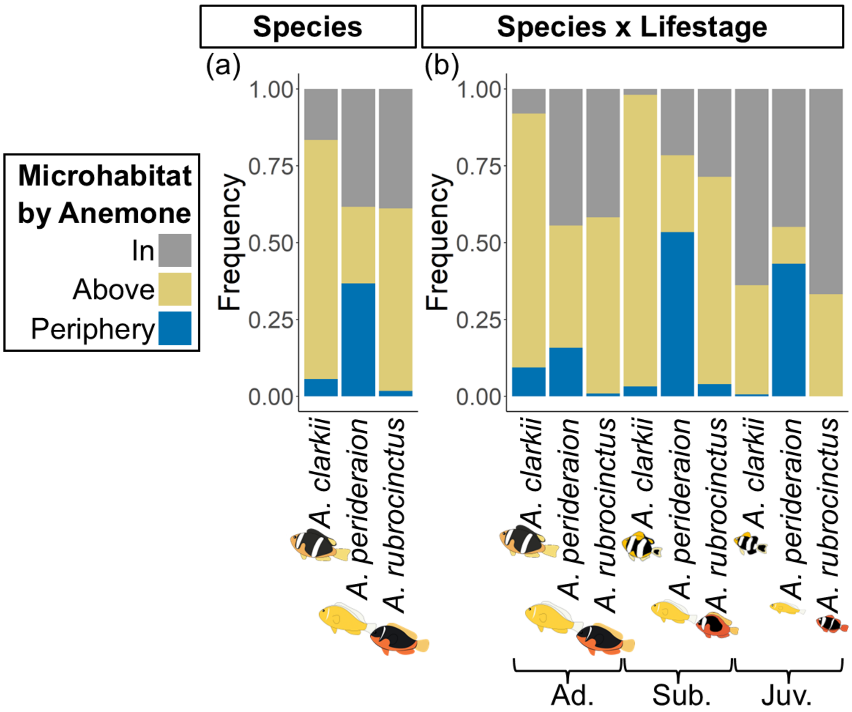


Supplementary Figure 2. Where the fish spent there time in relation to the microhabitat of the anemone.

*PREDATOR VISION AND CLOWNFISH COLOR-PATTERN DETECTABILITY*

**Overarching supplementary results:**

The mean and variance measurements were collected for each color pattern variable. The mean value represents a single score for all pixels within a clownfish image combined into a single average value and compared against the single value for the host anemone (i.e. brighter and duller regions of the clownfish image are averaged together into a single score). In contrast, the variance value captures pixel by pixel variation within each image and better reflect the larger amounts of variation produced by highly contrasting patterns (e.g. some parts of the body are far more variable than others in generalists with thick white stripes surrounded by black body color). Therefore, the variance value is included in the main text figure, and here is the mean value. *Amphiprion clarkii* generalists have higher mean luminance versus both specialist species compared to their anemone hosts ($p < 0.01$). The mean color (i.e. just noticeable difference) is lower for *A. clarkii* generalist than both specialist species compared to their anemone hosts ($p < 0.01$), yet this was due to black and white pixels being averaged together to make a gray mean body color.

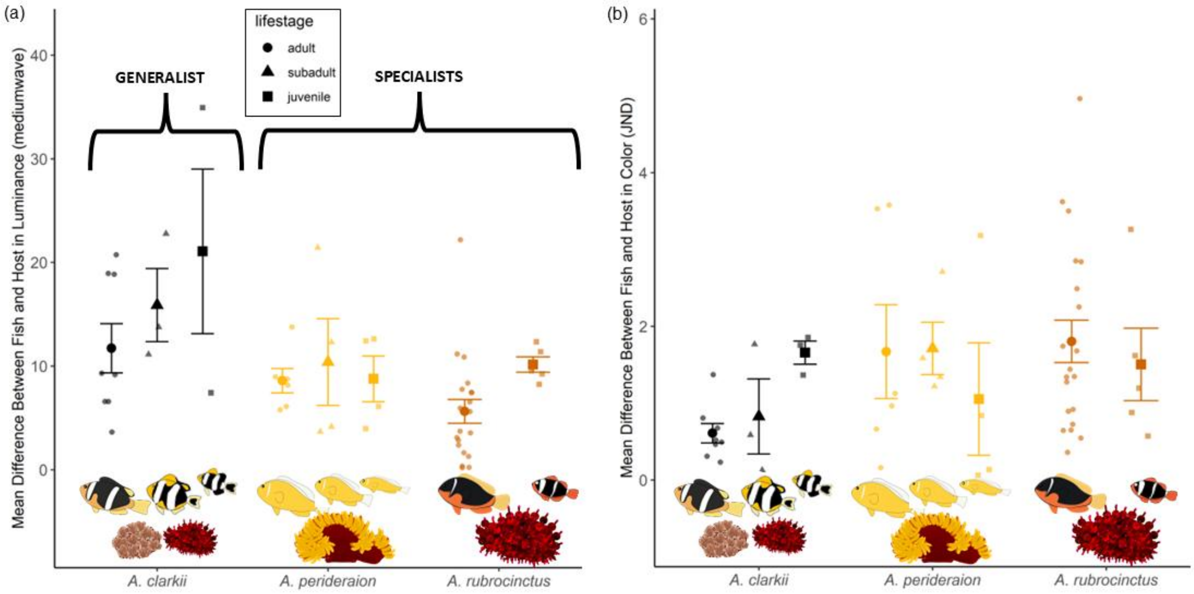


Supplementary Figure 3. Mean difference between fish and host in (a) luminance and (b) color pattern (just noticeable difference).

**Adults only: differences between fish and their own/same anemones, conspecific anemones, and other anemones**

Supplementary Table 2. Output of all statistical analyses for adult color patterns. LM = linear model; GLM = generalized linear model.

| Response Variable | Model | Predictor variable | Factor Type | df | Model Statistic | Test-value | p-value | R-squared |
|---|---|---|---|---|---|---|---|---|
| Diff between fish and hosts in | GLM | Species | fixed | 2 | χ2 (chi-squared) | 15.35 | 0.0005* | 0.36 |
| Mean Luminance 1m | | Host Type | fixed | 2 | χ2 (chi-squared) | 10.07 | 0.0065* | |
| (square-root transformed) | | Species x Host Type | interaction | 4 | χ2 (chi-squared) | 25.34 | < 0.0001* | |
| | | | | | Sample size | n = 102 | | |
| | | significant pairwise comparisons of interaction: | | | outliers removed | n = 2 | | |

| | | | | | | | | | |
|---|---|---|---|---|---|---|---|---|---|
| | | | | | | | A. clarkii own anemone vs A. rubrocinctus own anemone | 0.011* | |
| | | | | | | | A. rubrocinctus other anemone vs. A. rubrocinctus own anemone | 0.012* | |
| Diff between fish and hosts in | GLM | Species | fixed | 2 | $\chi2$ (chi-squared) | 32.33 | | < 0.0001* | 0.38 |
| Mean Color 1m | | Host Type | fixed | 2 | $\chi2$ (chi-squared) | 5.89 | | 0.0530 | |
| (fourth-root transformed) | | Species x Host Type | interaction | 4 | $\chi2$ (chi-squared) | 17.08 | | 0.002* | |
| Color = euclidean distance of diff. | | | | | Sample size | n = 102 | | | |
| color channels | | significant pairwise comparisons of interaction: | | | outliers removed | n = 1 | | | |
| | | | | | | | A. clarkii own anemone vs A. rubrocinctus own anemone | 0.012* | |
| | | | | | | | A. clarkii own anemone vs A. rubrocinctus other anemone | 0.0005* | |
| Diff between fish and hosts in | GLM | Species | fixed | 2 | $\chi2$ (chi-squared) | 101.47 | | < 0.0001* | 0.33 |
| Luminance Pattern Energy | | Distance | fixed | 2 | $\chi2$ (chi-squared) | 1.55 | | 0.46 | |
| a.k.a. variance of luminance | | Host Type | fixed | 2 | $\chi2$ (chi-squared) | 19.30 | | < 0.0001* | |
| (square-root transformed) | | Species x Distance | interaction | 4 | $\chi2$ (chi-squared) | 0.18 | | 1.00 | |
| | | Species x Host Type | interaction | 4 | $\chi2$ (chi-squared) | 11.38 | | 0.02* | |
| | | Distance x Host Type | interaction | 4 | $\chi2$ (chi-squared) | 0.50 | | 0.97 | |
| | | Species x Distance x Host Type | interaction | 8 | $\chi2$ (chi-squared) | 1.18 | | 1.00 | |
| | | | | | Sample size | n = 306 | | | |
| | | significant pairwise comparisons of interaction: | | | outliers removed | n = 1 | | | |
| | | | | | | | A. clarkii own anemone vs. all other species with each host type | <0.011* | |
| | | | | | | | A. clarkii other anemone vs A. rubrocinctus other anemone | 0.002* | |
| | | | | | | | A. clarkii other anemone vs. A. perideraion other anemone | 0.016* | |
| | | | | | | | A. rubrocinctus own anemone vs A. rubrocinctus other anemone | 0.01* | |
| Diff between fish and hosts in | GLM | Species | fixed | 2 | $\chi2$ (chi-squared) | 97.57 | | < 0.0001* | 0.31 |
| Color Pattern Energy | | Distance | fixed | 2 | $\chi2$ (chi-squared) | 6.07 | | 0.048* | |
| a.k.a. variance of color | | Host Type | fixed | 2 | $\chi2$ (chi-squared) | 2.03 | | 0.36 | |
| (square-root transformed) | | Species x Distance | interaction | 4 | $\chi2$ (chi-squared) | 10.42 | | 0.034* | |
| Color = euclidean distance of diff. | | Species x Host Type | interaction | 4 | $\chi2$ (chi-squared) | 9.17 | | 0.06 | |
| color channels | | Distance x Host Type | interaction | 4 | $\chi2$ (chi-squared) | 0.31 | | 0.99 | |
| | | Species x Distance x Host Type | interaction | 8 | $\chi2$ (chi-squared) | 0.47 | | 1.00 | |
| | | | | | Sample size | n = 306 | | | |
| | | | | | outliers removed | n = 3 | | | |
| | | significant pairwise comparisons of interaction: | | | | | | | |
| | | | | | | | A. clarkii 0.4m vs. other species distance combos | <0.002* | |
| | | | | | | | A. perideraion 0.4m vs. other species distance combos | <0.003* | |
| | | | | | | | A. rubrocinctus 0.4m vs. other species distance combos except vs. A. clarkii 5m | <0.036 | |
| | | | | | | | A. clarkii 1m vs. other species distance combos | <0.0008* | |
| | | | | | | | A. rubrocinctus 1m vs A. perideraion 1m | 0.003* | |
| | | | | | | | A. rubrocinctus 1m vs. A. perideraion 5m | 0.014* | |
| | | | | | | | A. rubrocinctus 5m vs. A. perideraion 1m | 0.035* | |
| | | | | | | | A. clarkii 5m vs. A. perideraion 5m | 0.032* | |
| | | | | | | | A. clarkii 0.4m vs. A. clarkii 5m | 0.035* | |
| | | | | | | | A. clarkii 1m vs. A clarkii 5m | 0.021* | |
| All fish and hosts separate | LM | Species | fixed | 5 | $\chi2$ (chi-squared) | 17.16 | | <0.0001* | 0.61 |
| Mean Luminance 1m | | | | | Sample size | n = 62 | | | |
| (log transformed) | | | | | outliers removed | n = 1 | | | |
| | | | significant pairwise comparisons of interaction: | | | | | | |
| | | | | | | | A. clarkii vs. A. rubrocinctus | <0.0001* | |
| | | | | | | | A. clarkii vs. E. quadricolor | <0.0001* | |
| | | | | | | | A. perideraion vs. A. rubrocinctus | 0.0009* | |
| | | | | | | | A. perideraion vs. E. quadricolor | 0.003* | |
| | | | | | | | A. rubrocinctus vs. H. crispa | 0.011* | |
| | | | | | | | E. quadricolor vs. H. crispa | 0.023* | |
| All fish and hosts separate | LM | Species | fixed | 5 | $\chi2$ (chi-squared) | 16.76 | | <0.0001* | 0.60 |
| Variance Luminance 1m | | | | | Sample size | n = 62 | | | |
| (log transformed) | | | | | outliers removed | n = 1 | | | |
| | | | significant pairwise comparisons of interaction: | | | | | | |
| | | | | | | | A. clarkii vs. all other species except H. crispa | <0.006* | |
| | | | | | | | A. perideraion vs. E. quadricolor | 0.010* | |
| | | | | | | | A. rubrocinctus vs. E. quadricolor | 0.0006* | |
| | | | | | | | E. quadricolor vs. H. crispa | 0.015* | |
| All fish and hosts separate | LM | Species | fixed | 5 | $\chi2$ (chi-squared) | 4.72 | | 0.001* | 0.30 |

| | | | | | | | | | |
|---|---|---|---|---|---|---|---|---|---|
| Mean Color 1m | | | | | Sample size | n = 62 | | | |
| (fourth-root transformed) | | | | | outliers removed | n = 0 | | | |
| Color = euclidean distance of diff. color channels | | | significant pairwise comparisons of interaction: | | | | A. clarkii vs. all other species except H. crispa | <0.039* | |
| All fish and hosts separate | LM | Species | fixed | 5 | $\chi2$ (chi-squared) | 19.78 | | <0.0001* | 0.64 |
| Variance in Color 1m | | | | | Sample size | n = 62 | | | |
| (fourth-root transformed) | | | | | outliers removed | n = 0 | | | |
| Color = euclidean distance of diff. color channels | | | significant pairwise comparisons of interaction: | | | | A. clarkii vs. all other species | <0.026* | |
| | | | | | | | A. perideraion vs. E. quadricolor | 0.002* | |
| | | | | | | | A. rubrocinctus vs. E. quadricolor | <0.0001* | |
| All fish and hosts separate | LM | Species | fixed | 5 | $\chi2$ (chi-squared) | 3.70 | | 0.006* | 0.26 |
| Mean X channel - Color 1m | | | | | Sample size | n = 62 | | | |
| (fourth-root transformed) | | | | | outliers removed | n = 2 | | | |
| X is difference in longwave and mediumwave | | | significant pairwise comparisons of interaction: | | | | A. clarkii vs. A. perideraion | 0.019* | |
| | | | | | | | A. clarkii vs. A. rubrocinctus | 0.018* | |
| | | | | | | | A. clarkii vs. H. magnifica | 0.040* | |
| All fish and hosts separate | LM | Species | fixed | 5 | $\chi2$ (chi-squared) | 43.33 | | <0.0001* | 0.80 |
| Variance X channel - Color 1m | | | | | Sample size | n = 62 | | | |
| (fourth-root transformed) | | | | | outliers removed | n = 1 | | | |
| X is difference in longwave and mediumwave | | | significant pairwise comparisons of interaction: | | | | A. clarkii vs. E. quadricolor | <0.0001* | |
| | | | | | | | A. clarkii vs. H. crispa | 0.002* | |
| | | | | | | | A. clarkii vs. H. magnifica | 0.009* | |
| | | | | | | | A. perideraion vs. A. rubrocinctus | 0.001* | |
| | | | | | | | A. perideraion vs. E .quadricolor | 0.0001* | |
| | | | | | | | A. rubrocinctus vs. E. quadricolor | <0.0001* | |
| | | | | | | | A. rubrocinctus vs. H. crispa | <0.0001* | |
| | | | | | | | A. rubrocinctus vs. H. magnifica | <0.0001* | |
| All fish and hosts separate | LM | Species | fixed | 5 | $\chi2$ (chi-squared) | 3.07 | | 0.016* | 0.22 |
| Mean Y channel - Color 1m | | | | | Sample size | n = 62 | | | |
| (fourth-root transformed) | | | | | outliers removed | n = 1 | | | |
| Y is difference between longwave & mediumwave with shortwave | | | significant pairwise comparisons of interaction: | | | | A. clarkii vs. H. magnifica | 0.019* | |
| All fish and hosts separate | LM | Species | fixed | 5 | $\chi2$ (chi-squared) | 15.38 | | <0.0001* | 0.58 |
| Variance Y channel - Color 1m | | | | | Sample size | n = 62 | | | |
| (fourth-root transformed) | | | | | outliers removed | n = 0 | | | |
| Y is difference between longwave & mediumwave with shortwave | | | significant pairwise comparisons of interaction: | | | | A. clarkii vs. A. rubrocinctus | 0.006* | |
| | | | | | | | A. clarkii vs. E. quadricolor | <0.0001* | |
| | | | | | | | A. clarkii vs. H. crispa | 0.01* | |
| | | | | | | | A. perideraion vs. E .quadricolor | 0.003* | |
| | | | | | | | A. rubrocinctus vs. E. quadricolor | <0.0001* | |

**Overall results:** The generalist was more luminant against all hosts than both specialists against their hosts ($p < 0.001$). At all distances, the pattern energy of luminance (i.e. variation) for generalists against all hosts was higher than both specialists ($p = 0.02$). The average color of generalists was less different to hosts than specialists ($p = 0.002$). However, it is important to note that generalist colors are black and white to make an average color of gray, which is less different to other colors. When predators view their prey, their eyes do not simply average the whole color of the fish, but instead they take into account the differentiation between color patterns on every part of the fish to find the location of the fish at striking distances. Accordingly, the pattern energy of color (i.e. variation in color pattern throughout the fish) was more different in generalists against all hosts compared to specialists at both 0.4m and 1m distance ($p < 0.034$). At a 5m distance, the generalist was more distinct in color pattern energy than one specialist (*A. perideraion*).

*Average luminance match (mediumwave): mean was square-root transformed*
All completed for 1m distance only.

- Mean luminance differences: The interaction between species and hosts was significant ($p < 0.001$). posthoc p-values below 0.05:
    - *A. clarkii* against its own host had higher values than *A. rubrocinctus* against its own host
    - *A. rubrocinctus* against other hosts had higher values than *A. rubrocinctus* against same hosts

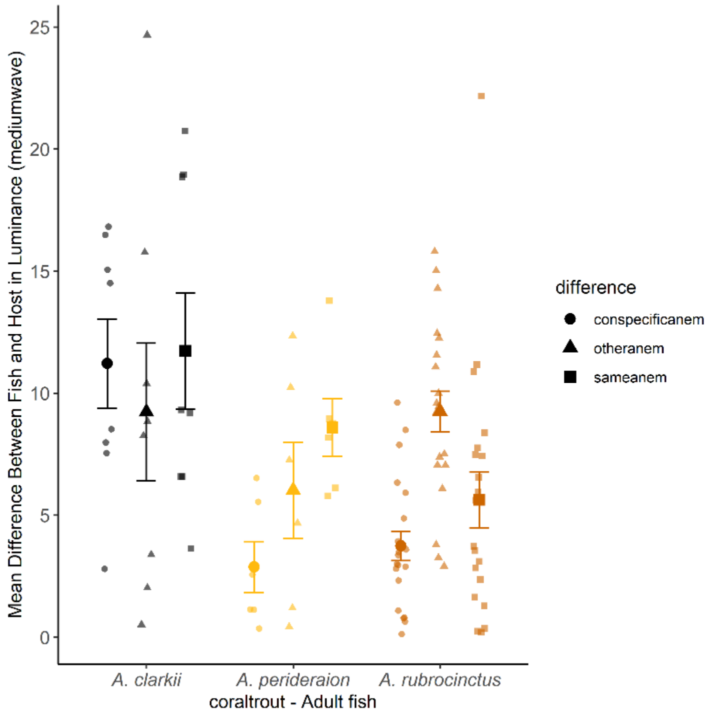


Supplementary Figure 4. Mean differences between adult fish and host in luminance for coral trout vision.

*Average color match (JND): mean was fourth-root transformed*

All completed for 1m distance only.

- Mean color differences: There was an interaction between species and type of host ($p = 0.002$). posthoc p-values below 0.05:
    - *A. clarkii* against its own host had lower mean color than *A. rubrocinctus* against its own host.
    - *A. clarkii* against its own host had lower mean color than *A. rubrocinctus* against other hosts.

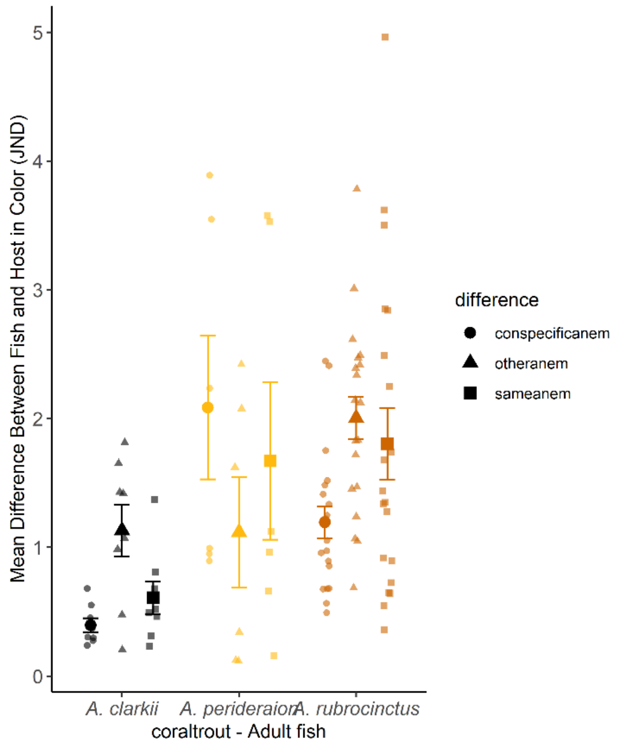


Supplementary Figure 5. Mean difference between adult fish and host for color (JND = just noticeable difference) for coral trout vision.

*Luminance pattern energy (mediumwave): variance was square-root transformed*
Used following distances: 0.4m, 1m, 5m

- Variance luminance differences: There was an interaction between species and types of hosts ($p = 0.02$). posthoc p-values below 0.05:
    - *A. clarkii* against its own host had higher luminance pattern energy than all other clownfish and host types.
    - *A. clarkii* against other host species was higher than *A. rubrocinctus* and *A. perideraion* against other host species.
    - *A. rubrocinctus* against its own host was higher in pattern energy than *A. rubrocinctus* against other host species.

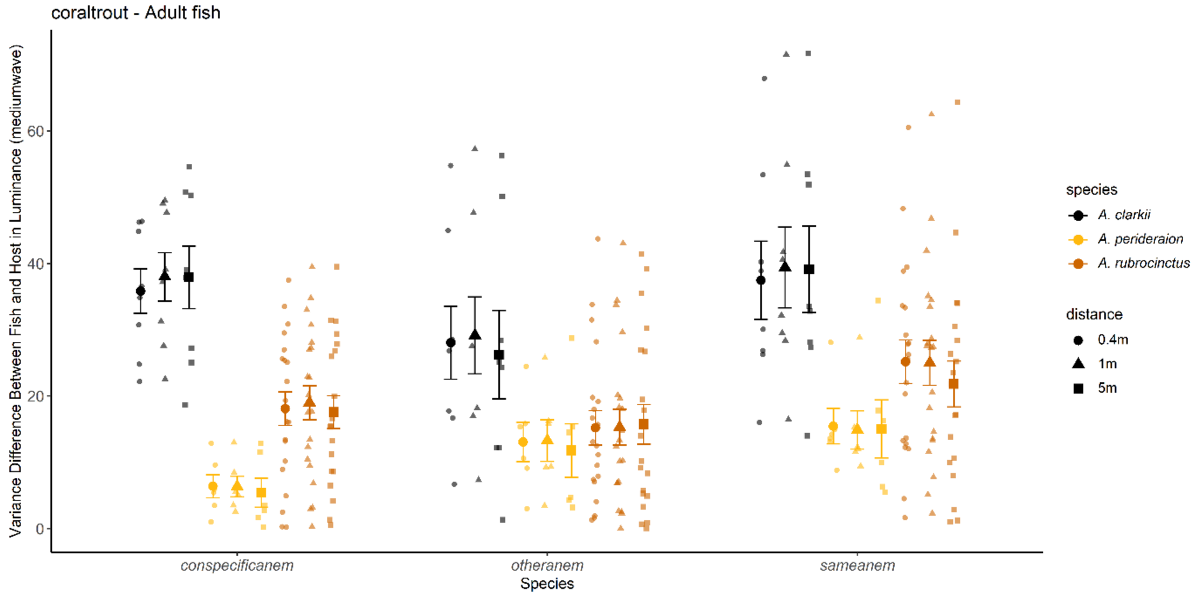

Supplementary Figure 6. Variance difference between adult fish and host for luminance for coral trout vision.

*Color pattern energy (JND): variance was fourth-root transformed*
Used following distances: 0.4m, 1m, 5m

- Variance color differences: There were interactions between species and distance ($p < 0.034$). posthoc p-values below 0.05:
    - Species x Distance
        - *A. clarkii* at 0.4m distance had higher color pattern energy than all other species at different distances. *A. clarkii* at 0.4m distance and 1m had higher color pattern energy than *A. clarkii* at 5m.
        - *A. perideraion* at 0.4m distance had lower color pattern energy than all other species at different distances.
        - *A. rubrocinctus* at 0.4m distance had that fell between all the other species at different distances.
        - *A. clarkii* at 1m distance had higher color pattern energy than all other species at different distances.
        - *A. rubrocinctus* at 1m distance had higher color pattern energy than *A. perideraion* at both 1m and 5m.
        - *A. clarkii* at 5m had higher color pattern energy than *A. perideraion* at 5m.

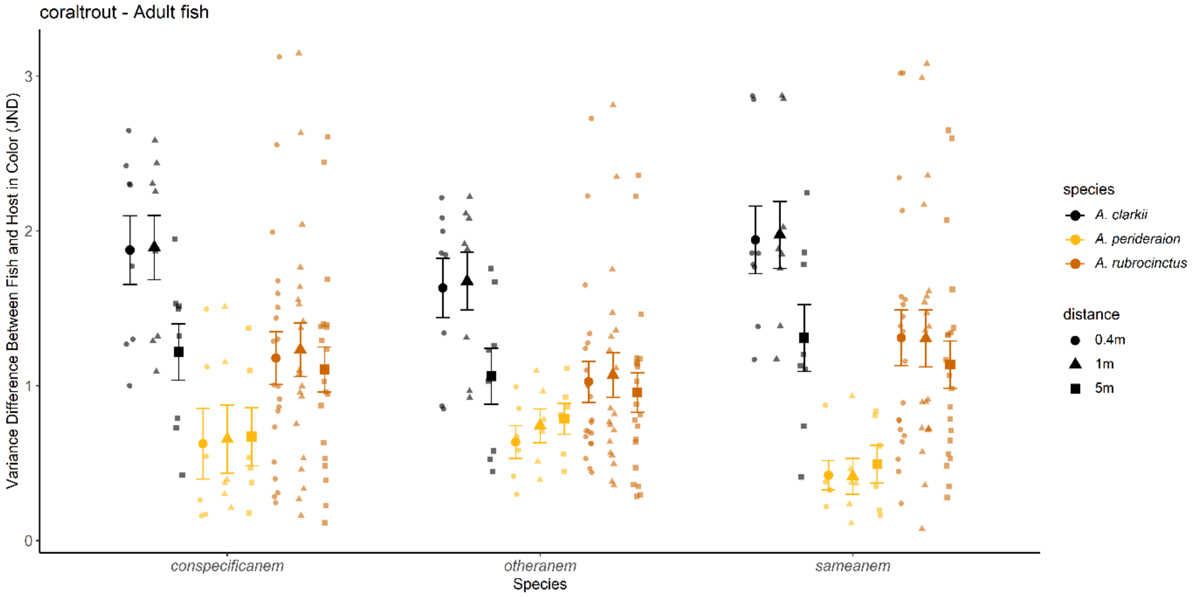


Supplementary Figure 7. Variance difference between adult fish and host for color (JND = just noticeable difference) for coral trout vision.

#### Subadults only: Differences between fish and their own/same anemones, conspecific anemones, and other anemones

Supplementary Table 3. Output of all statistical analyses for subadults color patterns. LM = linear model; GLM = generalized linear model.

| Response Variable | Model | Predictor variable | Factor Type | df | Model Statistic | Test-value | p-value | R-squared |
|---|---|---|---|---|---|---|---|---|
| Diff between fish and hosts in | GLM | Species | fixed | 1 | χ2 (chi-squared) | 2.40 | 0.12 | 0.26 |
| Mean Luminance 1m | | Host Type | fixed | 2 | χ2 (chi-squared) | 0.09 | 0.96 | |

| | | | | | | | | | |
|---|---|---|---|---|---|---|---|---|---|
| (square-root transformed) | | Species x Host Type | interaction | 2 | $\chi$2 (chi-squared) | 2.80 | | 0.25 | |
| | | | | | Sample size | n = 21 | | | |
| | | | | | outliers removed | n = 0 | | | |
| Diff between fish and hosts in | GLM | Species | fixed | 1 | $\chi$2 (chi-squared) | 7.33 | | 0.007* | 0.40 |
| Mean Color 1m | | Host Type | fixed | 2 | $\chi$2 (chi-squared) | 1.39 | | 0.50 | |
| (square-root transformed) | | Species x Host Type | interaction | 2 | $\chi$2 (chi-squared) | 1.38 | | 0.50 | |
| Color = euclidean distance of diff. | | | | | Sample size | n = 21 | | | |
| color channels | | significant pairwise comparisons of interaction: | | | outliers removed | n = 0 | | | |
| | | | | | | | A. clarkii own host vs. A. perideraion own host | 0.016* | |
| Diff between fish and hosts in | GLM | Species | fixed | 1 | $\chi$2 (chi-squared) | 89.50 | | <0.0001* | 0.75 |
| Luminance Pattern Energy | | Distance | fixed | 2 | $\chi$2 (chi-squared) | 3.67 | | 0.16 | |
| a.k.a. variance of luminance | | Host Type | fixed | 2 | $\chi$2 (chi-squared) | 11.28 | | 0.004* | |
| (square-root transformed) | | Species x Distance | interaction | 2 | $\chi$2 (chi-squared) | 0.24 | | 0.89 | |
| | | Species x Host Type | interaction | 2 | $\chi$2 (chi-squared) | 15.50 | | 0.0004* | |
| | | Distance x Host Type | interaction | 4 | $\chi$2 (chi-squared) | 3.84 | | 0.43 | |
| | | Species x Distance x Host Type | interaction | 4 | $\chi$2 (chi-squared) | 2.62 | | 0.62 | |
| | | | | | Sample size | n = 63 | | | |
| | | significant pairwise comparisons of interaction: | | | outliers removed | n = 1 | | | |
| | | | | | | | A. clarkii own host vs. A. perideraion all hosts | < 0.0001* | |
| | | | | | | | A. clarkii own host vs. A. clarkii other host | 0.001* | |
| Diff between fish and hosts in | GLM | Species | fixed | 1 | $\chi$2 (chi-squared) | 0.03 | | 0.87 | 0.45 |
| Color Pattern Energy | | Distance | fixed | 2 | $\chi$2 (chi-squared) | 18.05 | | 0.0001* | |
| a.k.a. variance of color | | Host Type | fixed | 2 | $\chi$2 (chi-squared) | 2.67 | | 0.26 | |
| (square-root transformed) | | Species x Distance | interaction | 2 | $\chi$2 (chi-squared) | 5.69 | | 0.058 | |
| Color = euclidean distance of diff. | | Species x Host Type | interaction | 2 | $\chi$2 (chi-squared) | 10.03 | | 0.007* | |
| color channels | | Distance x Host Type | interaction | 4 | $\chi$2 (chi-squared) | 0.49 | | 0.97 | |
| | | Species x Distance x Host Type | interaction | 4 | $\chi$2 (chi-squared) | 0.19 | | 1.00 | |
| | | | | | Sample size | n = 63 | | | |
| | | | | | outliers removed | n = 0 | | | |
| | | significant pairwise comparisons of interaction: | | | | | | | |
| | | | | | | | all values of 5m vs. 0.4m and 1m | <0.01* | |
| | | | | | | | A. clarkii own host vs. A. perideraion own host | < 0.01* | |
| All fish and hosts separate | LM | Species | fixed | 4 | $\chi$2 (chi-squared) | 13.42 | | <0.0001* | 0.63 |
| Mean Luminance 1m | | | | | Sample size | n = 37 | | | |
| (log transformed) | | | | | outliers removed | n = 1 | | | |
| | | | significant pairwise comparisons of interaction: | | | | | | |
| | | | | | | | A. clarkii vs. E. quadricolor | <0.0001* | |
| | | | | | | | A. clarkii vs. R. magnifica | 0.027* | |
| | | | | | | | A. perideraion vs. E. quadricolor | 0.0006* | |
| All fish and hosts separate | LM | Species | fixed | 4 | $\chi$2 (chi-squared) | 19.80 | | <0.0001* | 0.71 |
| Variance Luminance 1m | | | | | Sample size | n = 37 | | | |
| (log transformed) | | | | | outliers removed | n = 0 | | | |
| | | | significant pairwise comparisons of interaction: | | | | | | |
| | | | | | | | A. clarkii vs. E. quadricolor | <0.0001* | |
| | | | | | | | A. clarkii vs. R. magnifica | 0.002* | |
| | | | | | | | A. perideraion vs. E. quadricolor | 0.0006* | |
| | | | | | | | E. quadricolor vs. R. crispa | 0.003* | |
| All fish and hosts separate | LM | Species | fixed | 4 | $\chi$2 (chi-squared) | 5.36 | | 0.002* | 0.40 |
| Mean Color 1m | | | | | Sample size | n = 37 | | | |
| (fourth-root transformed) | | | | | outliers removed | n = 0 | | | |
| Color = euclidean distance of diff. | | | significant pairwise comparisons of interaction: | | | | | | |
| color channels | | | | | | | A. clarkii vs. A. perideraion and R. magnifica | <0.02* | |
| | | | | | | | A. perideraion vs. E. quadricolor | 0.025* | |
| All fish and hosts separate | LM | Species | fixed | 4 | $\chi$2 (chi-squared) | 12.17 | | <0.0001* | 0.60 |
| Variance in Color 1m | | | | | Sample size | n = 37 | | | |
| (square-root transformed) | | | | | outliers removed | n = 0 | | | |
| Color = euclidean distance of diff. | | | significant pairwise comparisons of interaction: | | | | | | |
| color channels | | | | | | | A. clarkii vs. E. quadricolor | 0.0002* | |
| | | | | | | | A. perideraion vs. E. quadricolor | 0.0001* | |
| All fish and hosts separate | LM | Species | fixed | 4 | $\chi$2 (chi-squared) | 3.92 | | 0.01* | 0.33 |
| Mean X channel - Color 1m | | | | | Sample size | n = 37 | | | |
| (square-root transformed) | | | | | outliers removed | n = 0 | | | |

| | | | | | | | | | |
|---|---|---|---|---|---|---|---|---|---|
| X is difference in longwave and mediumwave | | | significant pairwise comparisons of interaction: | | | | | | |
| | | | | | | | A. clarkii vs. A. perideraion | 0.034* | |
| | | | | | | | A. perideraion vs. E. quadricolor | 0.033* | |
| All fish and hosts separate Variance X channel - Color 1m | LM | Species | fixed | 4 | χ2 (chi-squared) | 10.67 | | <0.0001* | 0.57 |
| | | | | | Sample size | n = 37 | | | |
| (log transformed) | | | | | outliers removed | n = 0 | | | |
| X is difference in longwave and mediumwave | | | significant pairwise comparisons of interaction: | | | | | | |
| | | | | | | | A. clarkii vs. E. quadricolor | 0.003* | |
| | | | | | | | A. perideraion vs. E. quadricolor | <0.0001* | |
| | | | | | | | A. perideraion vs. R. crispa | 0.031* | |
| All fish and hosts separate | LM | Species | fixed | 4 | χ2 (chi-squared) | 3.70 | | 0.014* | 0.32 |
| Mean Y channel - Color 1m | | | | | Sample size | n = 37 | | | |
| (fourth-root transformed) | | | | | outliers removed | n = 1 | | | |
| Y is difference between longwave & mediumwave with shortwave | | | significant pairwise comparisons of interaction: | | | | | | |
| | | | | | | | A. clarkii vs. A. perideraion | 0.018* | |
| All fish and hosts separate Variance Y channel - Color 1m | LM | Species | fixed | 4 | χ2 (chi-squared) | 8.67 | | <0.0001* | 0.52 |
| | | | | | Sample size | n = 37 | | | |
| (log transformed) | | | | | outliers removed | n = 0 | | | |
| Y is difference between longwave & mediumwave with shortwave | | | significant pairwise comparisons of interaction: | | | | | | |
| | | | | | | | A. clarkii vs. E. quadricolor | 0.002* | |
| | | | | | | | A. perideraion vs. E. quadricolor | 0..001* | |

**Overall results:** Although the average luminance did not differ among clownfishes against their hosts (p > 0.12), the luminance pattern energy of the generalist was higher than the specialist at the three distances (p < 0.001). Although the average color was lower for generalists against their hosts than specialists ( p = 0.007), the average color of generalists would make gray. The color pattern energy was higher in generalist clownfish against their hosts than the specialist clownfish at 0.4m and 1m.

*Average luminance match (mediumwave): mean was square-root transformed*

All completed for 1m distance only.

- Mean luminance differences: No differences observed (p > 0.12).

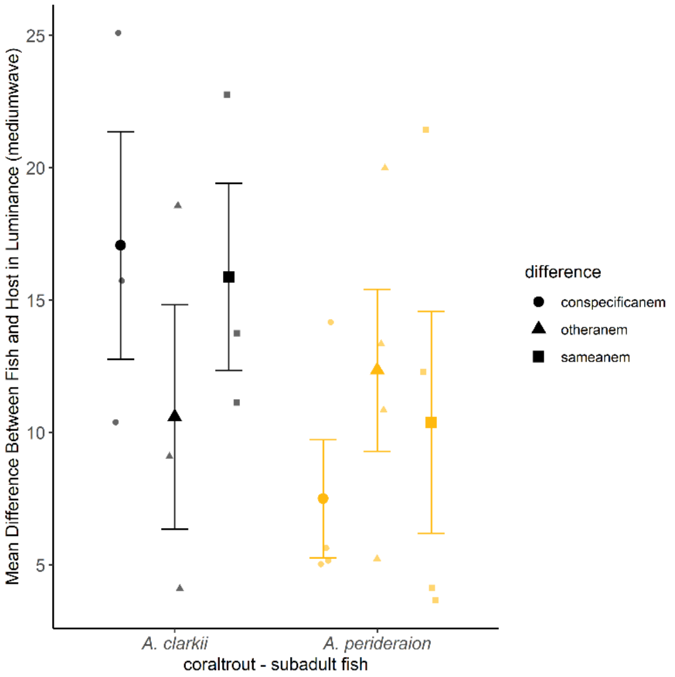


Supplementary Figure 8. mean difference between subadult fish and host for luminance for coral trout vision.

*Average color match (JND): mean was square-root transformed*

All completed for 1m distance only.

- Mean color differences: There were difference in species only (p = 0.007). posthoc p-values below 0.05:
    - *A. clarkii* against its own host had lower mean color than *A. perideraion* against its own host

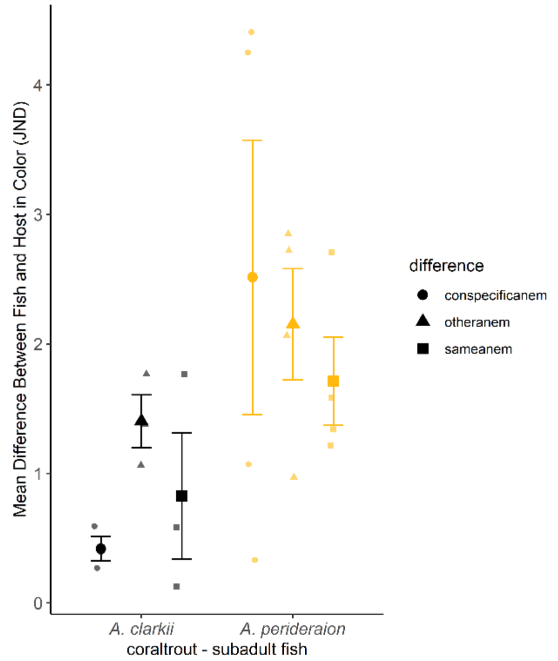


Supplementary Figure 9. Mean difference between subadult fish and host for color (JND = just noticeable difference) for coral trout vision.

*Luminance pattern energy (mediumwave): variance was square-root transformed*
Used following distances: 0.4m, 1m, 5m

- Variance luminance differences: There were differences in the interaction between species and host type (p < 0.001). posthoc p-values below 0.05:
    - *A. clarkii* against its own host was higher than *A. perideraion* on all hosts and than A. clarkii on other hosts

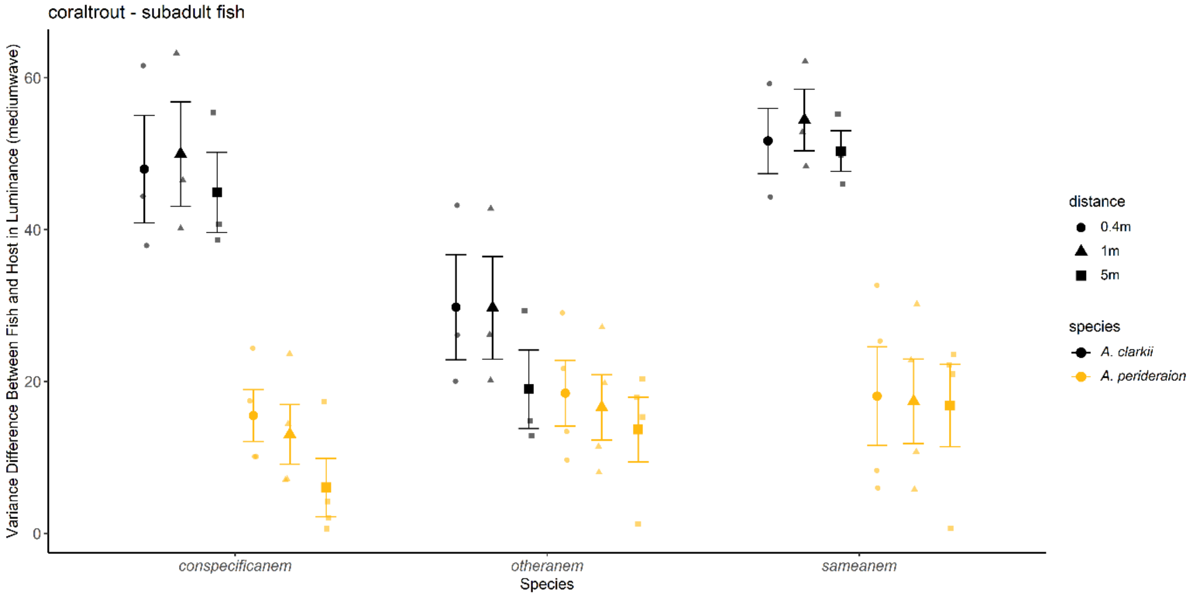

Supplementary Figure 10. Variance difference between subadult fish and host for luminance for coral trout vision.

*Color pattern energy (JND): variance was square-root transformed*
Used following distances: 0.4m, 1m, 5m

- Variance color differences: There were differences in distance ($p < 0.001$) and species ($p < 0.001$), but no interactions ($p = 0.06$). posthoc p-values below 0.05:
    - Species x Distance
        - All values at 5m were lower than other distances.
        - *A. clarkii* had higher color variance than *A. perideraion* on own host

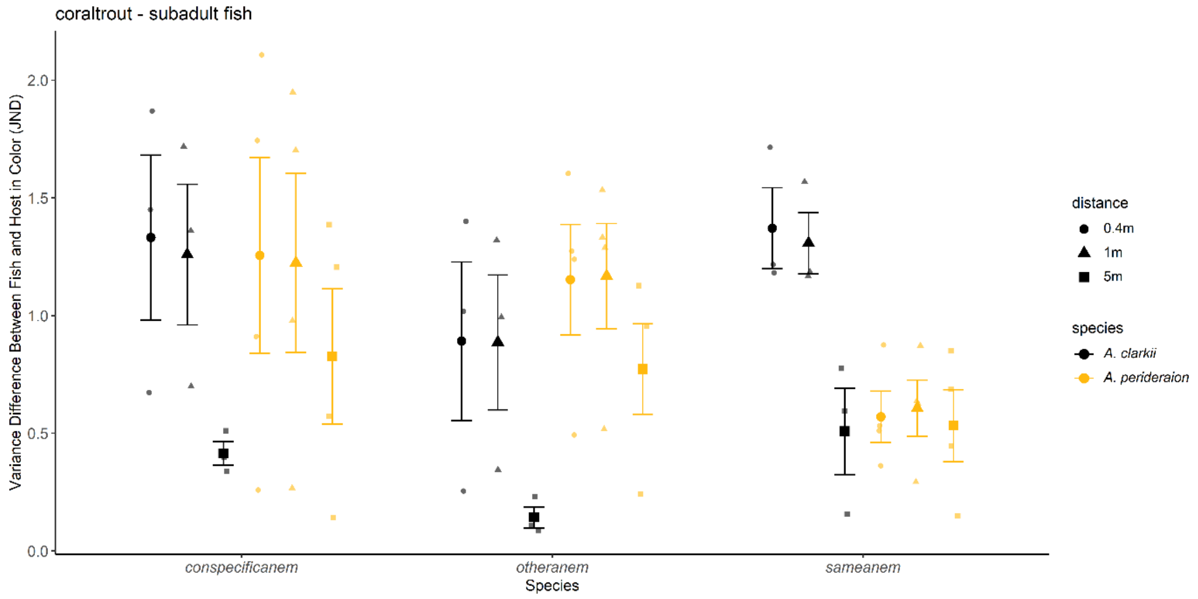


Supplementary Figure 11. Variance difference between subadult fish and host for color (JND = just noticeable difference) for coral trout vision.

#### Juveniles only: differences between fish and their own/same anemones, conspecific anemones, and other anemones

Supplementary Table 4. Output of all statistical analyses for juvenile color patterns. LM = linear model; GLM = generalized linear model.

| Response Variable | Model | Predictor variable | Factor Type | df | Model Statistic | Test-value | | p-value | R-squared |
|---|---|---|---|---|---|---|---|---|---|
| Diff between fish and hosts in | GLM | Species | fixed | 2 | χ2 (chi-squared) | 38.01 | | <0.0001* | 0.66 |
| Mean Luminance 1m | | Host Type | fixed | 2 | χ2 (chi-squared) | 1.59 | | 0.45 | |
| (square-root transformed) | | Species x Host Type | interaction | 4 | χ2 (chi-squared) | 11.04 | | 0.026* | |
| | | | | | Sample size | n = 36 | | | |
| | | significant pairwise comparisons of interaction: | | | outliers removed | n = 1 | | | |
| | | | | | | | A. clarkii own host vs. A. rubrocinctus other host | 0.012* | |
| | | | | | | | A. clarkii other host vs. A. rubrocinctus other host | 0.013* | |
| Diff between fish and hosts in | GLM | Species | fixed | 2 | χ2 (chi-squared) | 4.56 | | 0.10* | 0.47 |
| Mean Color 1m | | Host Type | fixed | 2 | χ2 (chi-squared) | 2.05 | | 0.36* | |
| (square-root transformed) | | Species x Host Type | interaction | 4 | χ2 (chi-squared) | 17.04 | | 0.002* | |
| Color = euclidean distance of diff. | | | | | Sample size | n = 36 | | | |

| | | | | | | | | | |
|---|---|---|---|---|---|---|---|---|---|
| color channels | | significant pairwise comparisons of interaction: | | | outliers removed | n = 1 | | | |
| | | | | | | | A. clarkii own host vs. A. perideraion own host | 0.018* | |
| Diff between fish and hosts in | GLM | Species | fixed | 2 | $\chi2$ (chi-squared) | 155.02 | | <0.0001* | 0.75 |
| Luminance Pattern Energy | | Distance | fixed | 2 | $\chi2$ (chi-squared) | 15.29 | | 0.0005* | |
| a.k.a. variance of luminance | | Host Type | fixed | 2 | $\chi2$ (chi-squared) | 11.91 | | 0.003* | |
| (square-root transformed) | | Species x Distance | interaction | 4 | $\chi2$ (chi-squared) | 4.41 | | 0.35 | |
| | | Species x Host Type | interaction | 4 | $\chi2$ (chi-squared) | 26.24 | | <0.0001* | |
| | | Distance x Host Type | interaction | 4 | $\chi2$ (chi-squared) | 3.15 | | 0.53 | |
| | | Species x Distance x Host Type | interaction | 8 | $\chi2$ (chi-squared) | 8.30 | | 0.41 | |
| | | | | | Sample size | n = 105 | | | |
| | | significant pairwise comparisons of interaction: | | | outliers removed | n = 5 | | | |
| | | | | | | | all values of 5m vs. 0.4m and 1m | <0.01* | |
| | | | | | | | A. clarkii own anemone vs. A. perideraion own anemone | <0.0001* | |
| | | | | | | | A. rubrocinctus own anemone vs. A. perideraion own anemone | <0.0001* | |
| | | | | | | | A. rubrocinctus own anemone vs. A. rubrocinctus other anemone | <0.0001* | |
| | | | | | | | A. clarkii own anemone vs. A. rubrocinctus other anemone | <0.0001* | |
| | | | | | | | A. rubrocinctus own anemone vs. A. perideraion other anemone | <0.0001* | |
| | | | | | | | A. clarkii own anemone vs. A. perideraion other anemone | <0.0001* | |
| | | | | | | | A. perideraion own anemone vs. A. clarkii other anemone | <0.0001* | |
| | | | | | | | A. clarkii other anemone vs, A. rubrocinctus other anemone | <0.0001* | |
| | | | | | | | A. clarkii other anemone vs. A. perideraion other anemone | <0.0001* | |
| Diff between fish and hosts in | GLM | Species | fixed | 2 | $\chi2$ (chi-squared) | 28.07 | | <0.0001* | 0.52 |
| Color Pattern Energy | | Distance | fixed | 2 | $\chi2$ (chi-squared) | 32.97 | | <0.0001* | |
| a.k.a. variance of color | | Host Type | fixed | 2 | $\chi2$ (chi-squared) | 1.51 | | 0.47 | |
| (fourth-root transformed) | | Species x Distance | interaction | 4 | $\chi2$ (chi-squared) | 9.08 | | 0.06 | |
| Color = euclidean distance of diff. | | Species x Host Type | interaction | 4 | $\chi2$ (chi-squared) | 4.17 | | 0.38 | |
| color channels | | Distance x Host Type | interaction | 4 | $\chi2$ (chi-squared) | 2.09 | | 0.72 | |
| | | Species x Distance x Host Type | interaction | 8 | $\chi2$ (chi-squared) | 5.67 | | 0.68 | |
| | | | | | Sample size | n = 105 | | | |
| | | | | | outliers removed | n = 2 | | | |
| | | significant pairwise comparisons of interaction: | | | | | | | |
| | | | | | | | all values of 5m vs. 0.4m and 1m | <0.01* | |
| | | | | | | | A. clarkii vs. A. perideraion | 0.012* | |
| | | | | | | | A. clarkii vs. A. rubrocinctus | 0.002* | |
| All fish and hosts separate | LM | Species | fixed | 5 | $\chi2$ (chi-squared) | 17.16 | | <0.0001* | 0.61 |
| Mean Luminance 1m | | | | | Sample size | n = 62 | | | |
| (log transformed) | | | | | outliers removed | n = 1 | | | |
| | | | significant pairwise comparisons of interaction: | | | | | | |
| | | | | | | | A. clarkii vs. A. rubrocinctus | <0.0001* | |
| | | | | | | | A. clarkii vs. E. quadricolor | <0.0001* | |
| | | | | | | | A. perideraion vs. A. rubrocinctus | 0.0009* | |
| | | | | | | | A. perideraion vs. E. quadricolor | 0.003* | |
| | | | | | | | A. rubrocinctus vs. H. crispa | 0.011* | |
| | | | | | | | E. quadricolor vs. H. crispa | 0.023* | |
| All fish and hosts separate | LM | Species | fixed | 5 | $\chi2$ (chi-squared) | 16.76 | | <0.0001* | 0.60 |
| Variance Luminance 1m | | | | | Sample size | n = 62 | | | |
| (log transformed) | | | | | outliers removed | n = 1 | | | |
| | | | significant pairwise comparisons of interaction: | | | | | | |
| | | | | | | | A. clarkii vs. all species except H. crispa | <0.007* | |
| | | | | | | | A. perideraion vs. E. quadricolor | 0.01* | |
| | | | | | | | A. rubrocinctus vs. E. quadricolor | 0.0006* | |
| | | | | | | | E. quadricolor vs. H. crispa | 0.015* | |
| All fish and hosts separate | LM | Species | fixed | 5 | $\chi2$ (chi-squared) | 5.25 | | 0.0005* | 0.32 |
| Mean Color 1m | | | | | Sample size | n = 62 | | | |
| (log transformed) | | | | | outliers removed | n = 0 | | | |
| Color = euclidean distance of diff. | | | significant pairwise comparisons of interaction: | | | | | | |
| color channels | | | | | | | A. clarkii vs. all other species except H. crispa | <0.014* | |
| All fish and hosts separate | LM | Species | fixed | 5 | $\chi2$ (chi-squared) | 19.78 | | <0.0001* | 0.64 |
| Variance in Color 1m | | | | | Sample size | n = 62 | | | |
| (fourth-root transformed) | | | | | outliers removed | n = 1 | | | |
| Color = euclidean distance of diff. | | | significant pairwise comparisons of interaction: | | | | | | |
| color channels | | | | | | | A. clarkii vs. all other species | <0.025* | |
| | | | | | | | A. perideraion vs. E. quadricolor | 0.002* | |
| | | | | | | | A. rubrocinctus vs. E. quadricolor | <0.0001* | |
| All fish and hosts separate | LM | Species | fixed | 5 | $\chi2$ (chi-squared) | 3.70 | | 0.006* | 0.26 |

| | | | | | | | | | |
|---|---|---|---|---|---|---|---|---|---|
| Mean X channel - Color 1m | | | | | Sample size | n = 62 | | | |
| (fourth-root transformed) | | | | | outliers removed | n = 2 | | | |
| X is difference in longwave and mediumwave | | | significant pairwise comparisons of interaction: | | | | | | |
| | | | | | | | A. clarkii vs. A. perideraion | 0.019* | |
| | | | | | | | A. clarkii vs. A. rubrocinctus | 0.018* | |
| | | | | | | | A. clarkii vs. H. magnifica | 0.04* | |
| All fish and hosts separate | LM | Species | fixed | 5 | χ2 (chi-squared) | 44.17 | | <0.0001* | 0.81 |
| Variance X channel - Color 1m | | | | | Sample size | n = 62 | | | |
| (square-root transformed) | | | | | outliers removed | n = 3 | | | |
| X is difference in longwave and mediumwave | | | significant pairwise comparisons of interaction: | | | | | | |
| | | | | | | | A. clarkii vs. all host species | <0.004* | |
| | | | | | | | A. perideraion vs. A. robrocinctus | 0.0005* | |
| | | | | | | | A. perideraion vs. E. quadricolor | 0.0001* | |
| | | | | | | | A. rubrocinctus vs. all host species | <0.0001* | |
| All fish and hosts separate | LM | Species | fixed | 5 | χ2 (chi-squared) | 3.07 | | 0.016* | 0.22 |
| Mean Y channel - Color 1m | | | | | Sample size | n = 62 | | | |
| (fourth-root transformed) | | | | | outliers removed | n = 1 | | | |
| Y is difference between longwave & mediumwave with shortwave | | | significant pairwise comparisons of interaction: | | | | | | |
| | | | | | | | A. clarkii vs. H. magnifica | 0.019* | |
| All fish and hosts separate | LM | Species | fixed | 5 | χ2 (chi-squared) | 18.01 | | <0.0001* | 0.62 |
| Variance Y channel - Color 1m | | | | | Sample size | n = 62 | | | |
| (fourth-root transformed) | | | | | outliers removed | n = 1 | | | |
| Y is difference between longwave & mediumwave with shortwave | | | significant pairwise comparisons of interaction: | | | | | | |
| | | | | | | | A. clarkii vs. all species | <0.016* | |
| | | | | | | | A. perideraion vs. E. quadricolor | 0.002* | |
| | | | | | | | A. rubrocinctus vs. E. quadricolor | <0.0001* | |

**Overall results:** The generalist clownfish was more luminant against its host than the specialist *A. rubrocinctus* ($p < 0.026$). The luminance pattern energy of the generalist was higher against hosts than both specialists clownfishes at all distances ($p < 0.001$). The generalist had higher average color against its host than both specialists ($p = 0.002$). The color pattern energy was higher on the generalist against its host than both specialists at all distances ($p < 0.001$) and the differences became a lot smaller at the further distance (5m).

*Average luminance match (mediumwave): mean was square-root transformed*
All completed for 1m distance only.

- Mean luminance differences: The interaction between species and hosts was significant ($p < 0.026$). posthoc p-values below 0.05:
  - *A. clarkii* against its own host had higher values than *A. rubrocinctus* against other hosts
  - *A. rubrocinctus* against other hosts had lower values than *A. rubrocinctus* against its own hosts

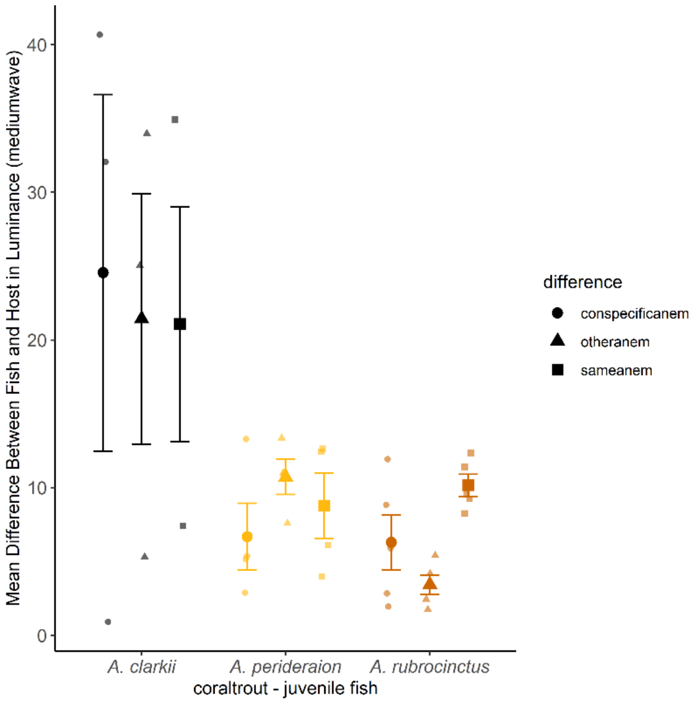

Supplementary Figure 12. Mean difference between juvenile fish and host for luminance for coral trout vision.

*Average color match (JND): mean was square-root transformed*

All completed for 1m distance only.

- Mean color differences: There was an interaction between species and type of host ($p = 0.002$). posthoc p-values below 0.05:
  - *A. clarkii* against its own host had higher mean color than *A. perideraion* against its own host

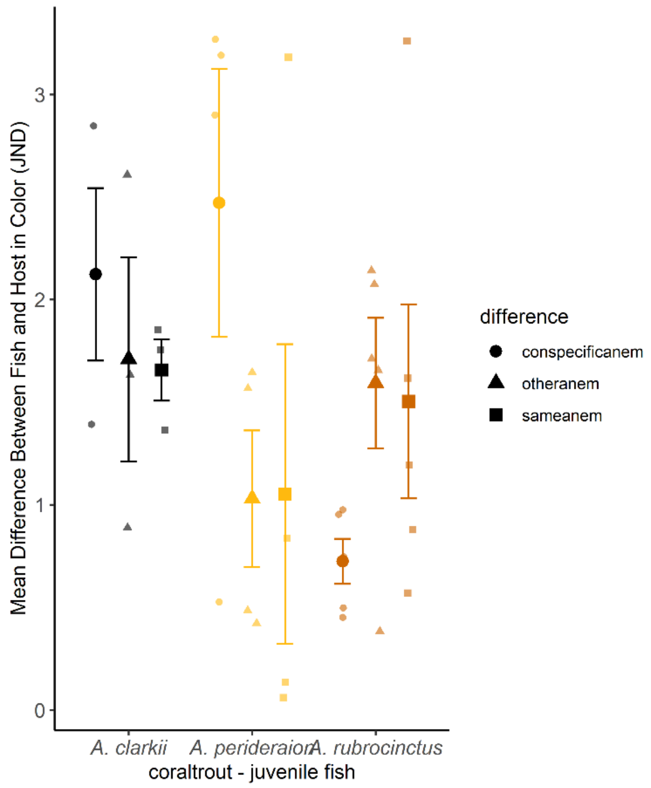


Supplementary Figure 13. Mean difference between juvenile fish and host for color (JND = just noticeable difference) for coral trout vision.

*Luminance pattern energy (mediumwave): variance was square-root transformed*
Used following distances: 0.4m, 1m, 5m

- Variance luminance differences: There were differences in the distance ($p = 0.0005$) without interaction with other factors. There was an interaction between species and types of hosts ($p < 0.001$). posthoc p-values below 0.05:
    - All differences at 5m were lower than differences at other distances.
    - *A. clarkii* against its own host was higher than *A. perideraion* against its own host and both specialist against other hosts
    - *A. rubrocinctus* against its own host was higher than *A. perideraion* against its own host and other hosts
    - *A. rubrocinctus* against its own host was higher than against other hosts.

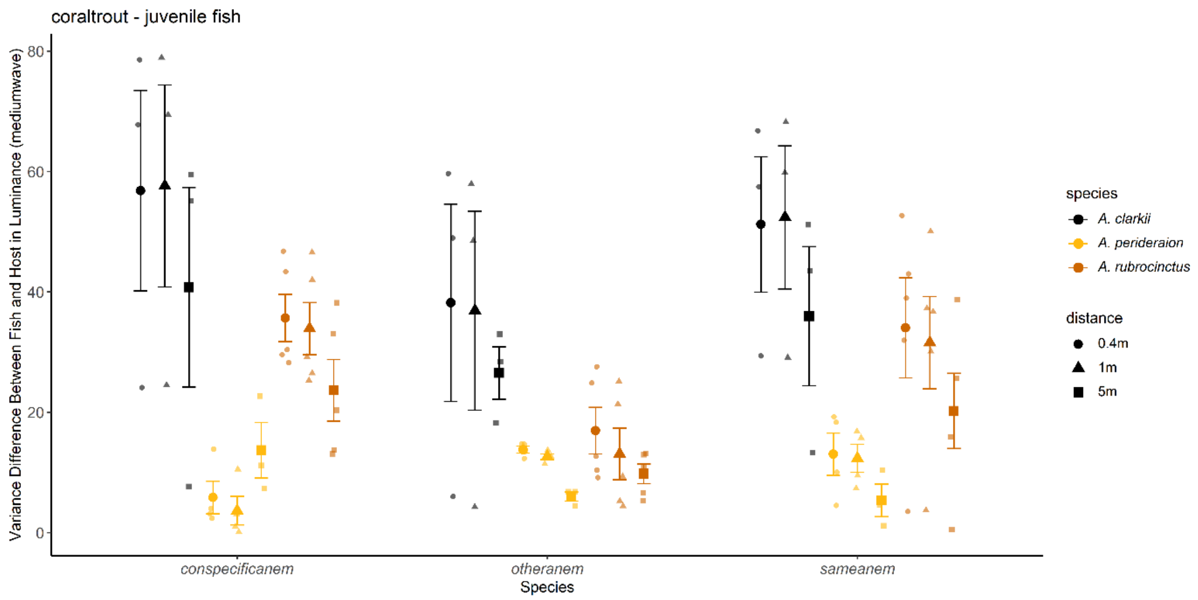


Supplementary Figure 14. Variance difference between juvenile fish and host for luminance for coral trout vision.

*Color pattern energy (JND): variance was fourth-root transformed*

Used following distances: 0.4m, 1m, 5m

- Variance color differences: There were differences in distance ($p < 0.001$) and species ($p < 0.001$), but no interactions ($p = 0.06$). posthoc p-values below 0.05:
    - Species x Distance
        - All values at 5m were lower than other distances.
        - *A. clarkii* had higher color variance than *A. perideraion* and *A. rubrocinctus*

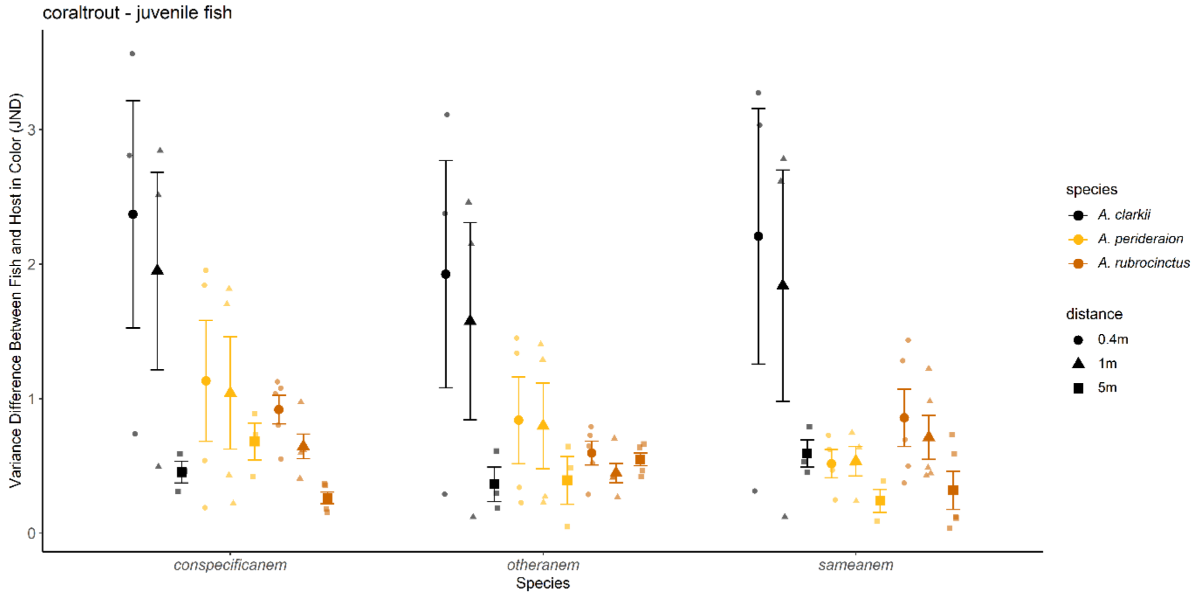


Supplementary Figure 15. Variance difference between juvenile fish and host for color (JND = just noticeable difference) for coral trout vision.

#### ***DIETARY NICHE***

Supplementary Table 5. Statistical output for all stable isotope analyses. LM = linear model.

| Response Variable | Model | Predictor variable | Factor Type | df | Model Statistic | Test-value | p-value | R-squared |
|---|---|---|---|---|---|---|---|---|
| Average Trophic Position | LM | Species | fixed | 4 | F-value | 400111.00 | < 0.0001* | 0.97 |
| Clownfish with Anemone pooled | | | | | Sample size | n = 48788 | | |
| and predatory fish pooled | | | | | Outliers removed | n = 1237 | | |
| square-root transformed | pairwise Tukey's | all pairwise interactions were significant to p < 0.0001* | | | | | | |
| Average delta Carbon | LM | Species | fixed | 4 | F-value | 13.14 | < 0.0001* | 0.51 |
| Clownfish with Anemone pooled | | | | | Sample size | n = 57 | | |
| and predatory fish pooled | | | | | Outliers removed | n = 1 | | |
| fourth-power transformed | pairwise | all pairwise interactions were not significant except: | | | | | | |
| | Tukey's | *Amphiprion clarkii - Amphiprion rubrocinctus* | | | t-ratio | 4.98 | < 0.0001* | |
| | | *Amphiprion clarkii - anemone* | | | | 2.83 | 0.0494* | |
| | | *Amphiprion perideraion - predatory fish* | | | | -4.13 | 0.0012* | |
| | | *Amphiprion rubrocinctus - predatory fish* | | | | -6.13 | < 0.0001* | |
| | | anemone - predatory fish | | | | -4.49 | 0.0004* | |
| Average Trophic Position | LM | Life stage | fixed | 1 | F-value | 13152.40 | < 0.0001* | 0.47 |
| Clownfish x Life stage | | Species | fixed | 2 | F-value | 11410.60 | < 0.0001* | |
| log10 transformed | | Life stage*Species | | 2 | F-value | 7636.60 | < 0.0001* | |
| | | | | | Sample size | n = 60030 | | |
| | pairwise | all pairwise interactions were significant to p < 0.0001* except: | | | Outliers removed | n = 2168 | | |
| | Tukey's | *adult Amphiprion clarkii - adult Amphiprion perideraion* | | | t-ratio | -0.53 | 0.9950 | |
| Average delta Carbon | LM | Life stage | fixed | 1 | F-value | 1.61 | 0.212 | 0.44 |
| Clownfish x Life stage | | Species | fixed | 2 | F-value | 9.19 | 0.0006* | |
| fourth-root transformed | | Life stage*Species | | 2 | F-value | 3.96 | 0.0280* | |
| | | | | | Sample size | n = 41 | | |
| | pairwise | all pairwise interactions were not significant except: | | | Outliers removed | n = 0 | | |
| | Tukey's | *adult Amphiprion clarkii - juvenile Amphiprion clarkii* | | | t-ratio | 3.25 | 0.0286* | |
| | | *adult Amphiprion clarkii - adult Amphiprion rubrocinctus* | | | | 4.86 | 0.0003* | |
| | | *adult Amphiprion clarkii - juvenile Amphiprion rubrocinctus* | | | | 4.10 | 0.0029* | |
| Average Trophic Position | LM | Region | fixed | 1 | F-value | 2932.15 | < 0.0001* | 0.88 |
| Adult Clownfish x Region | | Species | fixed | 1 | F-value | 287021.33 | < 0.0001* | |
| not transformed | | Region*Species | | 1 | F-value | 694.06 | < 0.0001* | |
| | pairwise | all pairwise interactions were significant to p < 0.0001* | | | Sample size | n = 40020 | | |
| | Tukey's | | | | Outliers removed | n = 676 | | |
| Average delta Carbon | LM | Region | fixed | 1 | F-value | 9.26 | 0.006* | 0.71 |
| Adult Clownfish x Region | | Species | | 1 | F-value | 39.59 | < 0.0001* | |
| log transformed | | Region*Species | | 1 | F-value | 4.61 | 0.0430* | |
| | | | | | Sample size | n = 27 | | |
| | pairwise | | | | Outliers removed | n = 1 | | |
| | Tukey's | *north Amphiprion clarkii - south Amphiprion clarkii* | | 22 | t-ratio | -2.96 | 0.0336* | |
| | | *north Amphiprion clarkii - north Amphiprion rubrocinctus* | | 22 | | 2.75 | 0.05338 | |
| | | *north Amphiprion clarkii - south Amphiprion rubrocinctus* | | 22 | | 2.13 | 0.17386 | |
| | | *south Amphiprion clarkii - north Amphiprion rubrocinctus* | | 22 | | 6.94 | < 0.0001* | |
| | | *south Amphiprion clarkii - south Amphiprion rubrocinctus* | | 22 | | 6.06 | < 0.0001* | |
| | | *north Amphiprion rubrocinctus - south Amphiprion rubrocinctus* | | 22 | | -0.74 | 0.87972 | |
| Average delta Nitrogen | paired t-test | | | 20 | t-value | 3.27 | 0.0038* | |
| Ethanol vs No Treatment | mean difference = 0.3429 | | | | sample size | n = 42 | | |
| Average delta Carbon | paired t-test | | | 20 | t-value | 0.62 | 0.5439 | |
| Ethanol vs No Treatment | mean difference = 0.0905 | | | | sample size | n = 42 | | |

Isotopic biplots show partitioning along the $\delta^{15}N$ axis and minimal partitioning along the $\delta^{13}C$ axis for clownfish compared to other species collected for reference. The predatory fish had considerably higher $\delta^{15}N$ than clownfish, which had higher $\delta^{15}N$ than most anemones and other invertebrates. However, compared to invertebrates and algae, clownfish $\delta^{13}C$ had limited variation and clustered together.

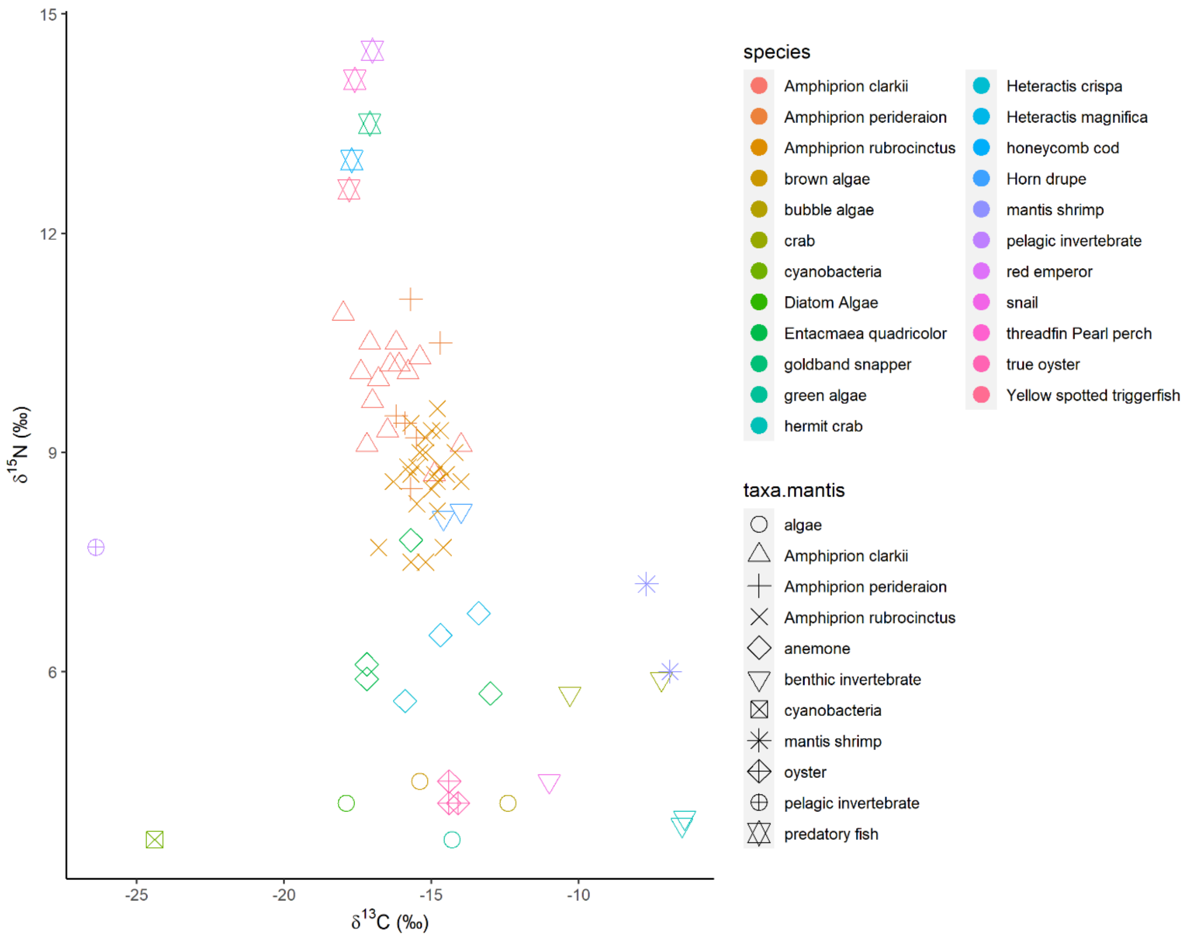


Supplementary Figure 16. Isotopic biplot of carbon and nitrogen values for all samples collected.

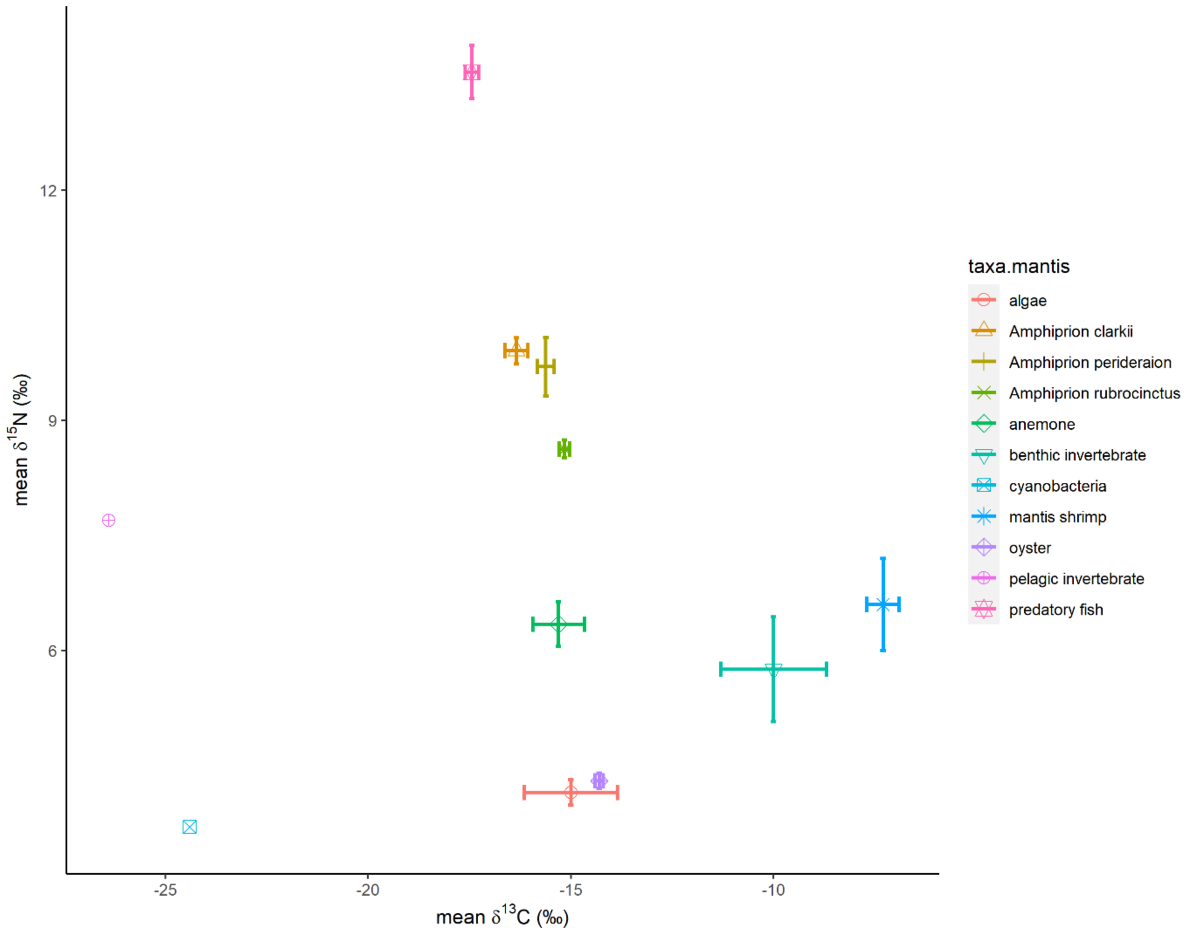


Supplementary Figure 17. Mean and standard error of carbon and nitrogen values in an isotopic biplot for all samples collected.

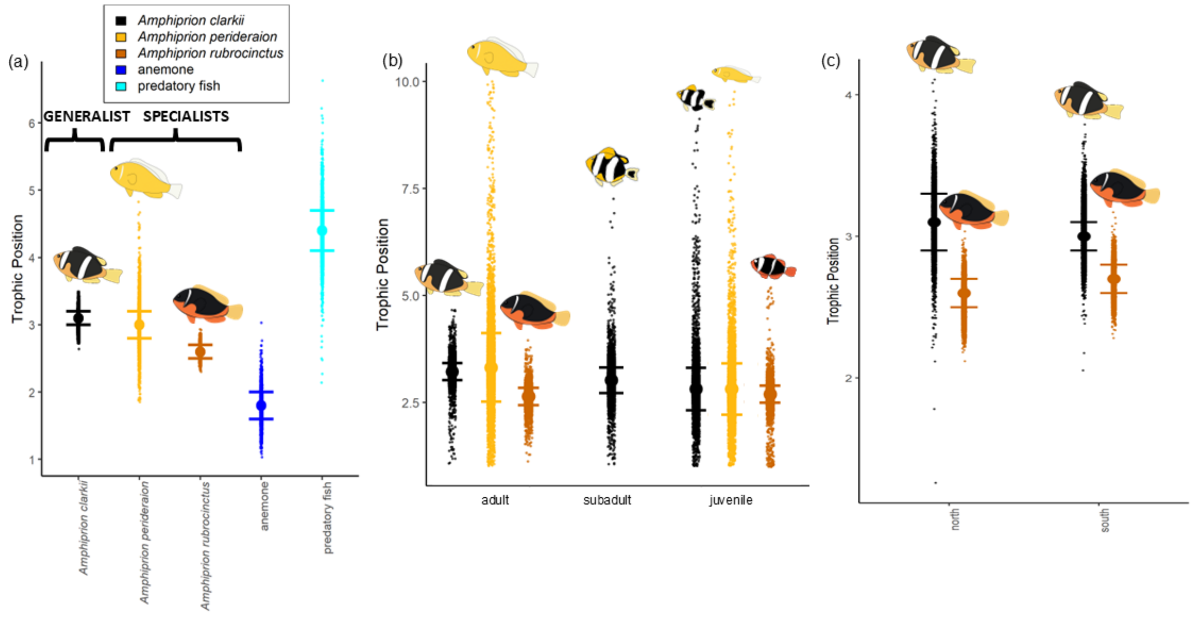


Supplementary Figure 18. Trophic positions of (a) clownfish species, pooled anemones, and pooled predatory fish; (b) clownfish life stages; and (c) clownfish by region (all *A.*

*perideraion* were located in the same region, therefore only regional comparison for the other two species).

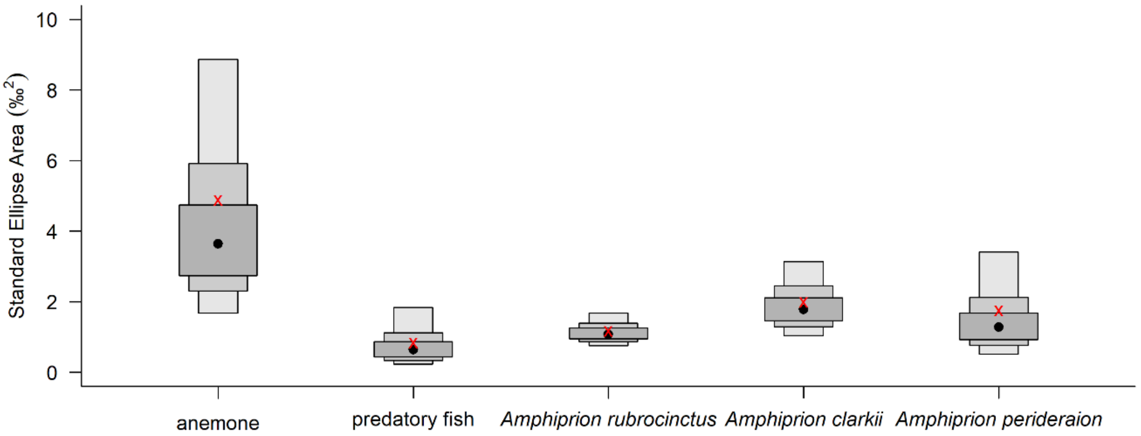


Supplementary Figure 19. Mean and standard error of standard ellipse areas for piscivorous fish, anemones, and clownfish species.

Supplementary Table 6. Isotopic niche overlap among piscivorous fish, clownfish species, and anemones for standard ellipses (40%).

**Overlap Niche Area (‰$^2$)** / **Percent Niche Overlap (%)**

| | *A.clarkii* | *A. perideraion* | *A. rubrocinctus* | anemone | predatory fish |
|---|---|---|---|---|---|
| *A.clarkii* | | 24% | <0.01% | <0.01% | <0.01% |
| *A. perideraion* | 0.74 | | 3% | <0.01% | 0% |
| *A. rubrocinctus* | <0.01 | 0.09 | | <0.01% | 0% |
| anemone | <0.01 | <0.01 | <0.01 | | <0.01% |
| predatory fish | <0.01 | 0 | 0 | <0.01 | |

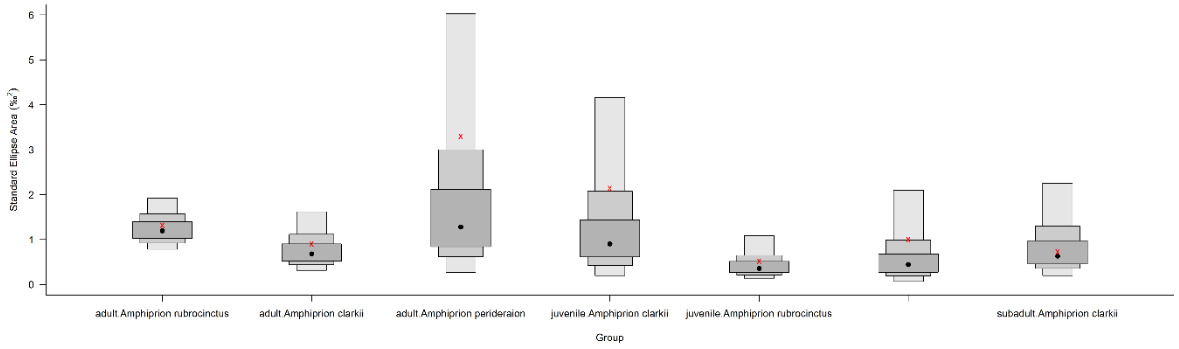

Supplementary Figure 20. Mean and standard error of standard ellipse areas for clownfish at different life stages.

Supplementary Table 7. Isotopic niche overlap among different life stages of clownfish species for standard ellipses (40%).

Percent Niche Overlap (%) (upper triangle); Overlap Niche Area (‰$^2$) (lower triangle)

| | *adult-A. clarkii* | *adult-A. perideraion* | *adult-A. rubrocinctus* | *subadult-A. clarkii* | *juvenile-A. clarkii* | *juvenile-A. perideraion* | *juvenile-A. rubrocinctus* |
|---|---|---|---|---|---|---|---|
| *adult-A. clarkii* | | 5% | <0.01% | 9% | <0.01% | <0.01% | 0% |
| *adult-A. perideraion* | 0.19 | | <0.01% | 7% | 6% | 10% | <0.01% |
| *adult-A. rubrocinctus* | <0.01 | <0.01 | | <0.01% | 18% | 16% | 29% |
| *subadult-A. clarkii* | 0.14 | 0.27 | <0.01 | | 0.32% | 0.15% | <0.01% |
| *juvenile-A. clarkii* | <0.01 | 0.29 | 0.54 | 0.01 | | 6% | 22% |
| *juvenile-A. perideraion* | <0.01 | 0.40 | 0.32 | 0.00 | 0.62 | | 9% |
| *juvenile-A. rubrocinctus* | 0.00 | <0.01 | 0.43 | <0.01 | 0.49 | 0.13 | |

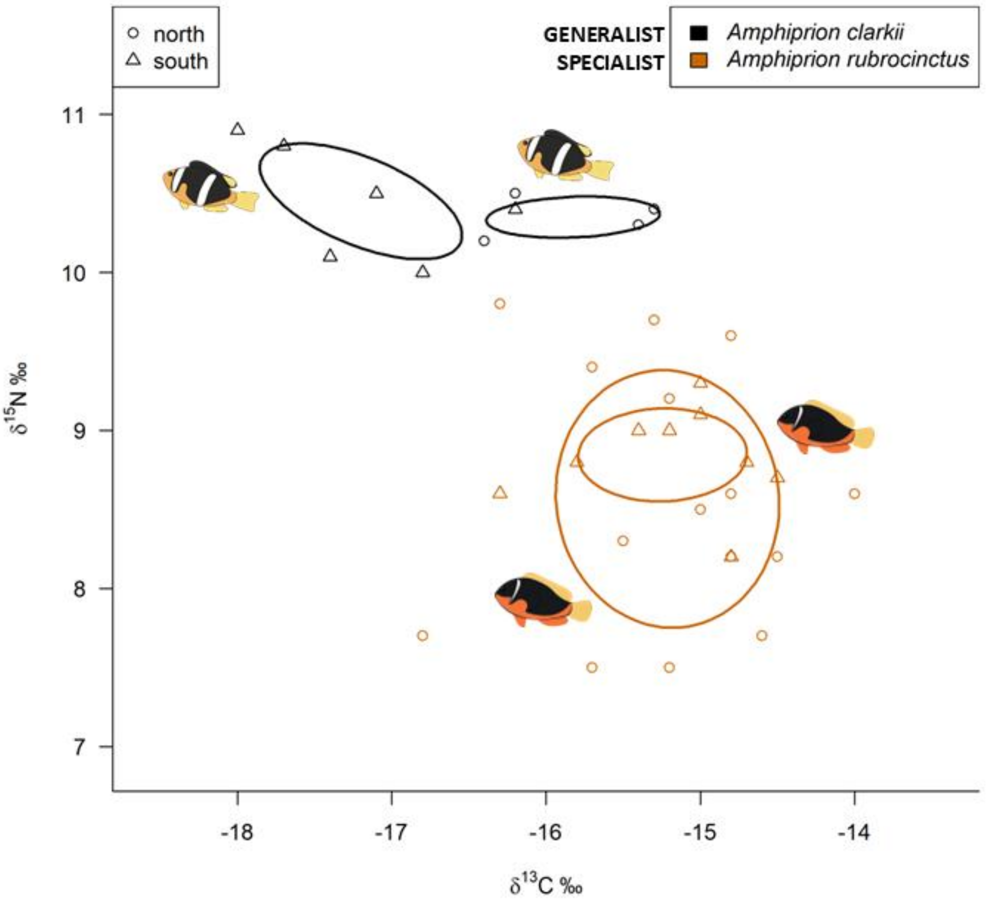


Supplementary Figure 21. Dietary niche overlap of clownfish by region (all *A. perideraion* were located in the same region, therefore only regional comparison for the other two species). Standard ellipses areas (40%) are depicted for carbon and nitrogen isotopes.

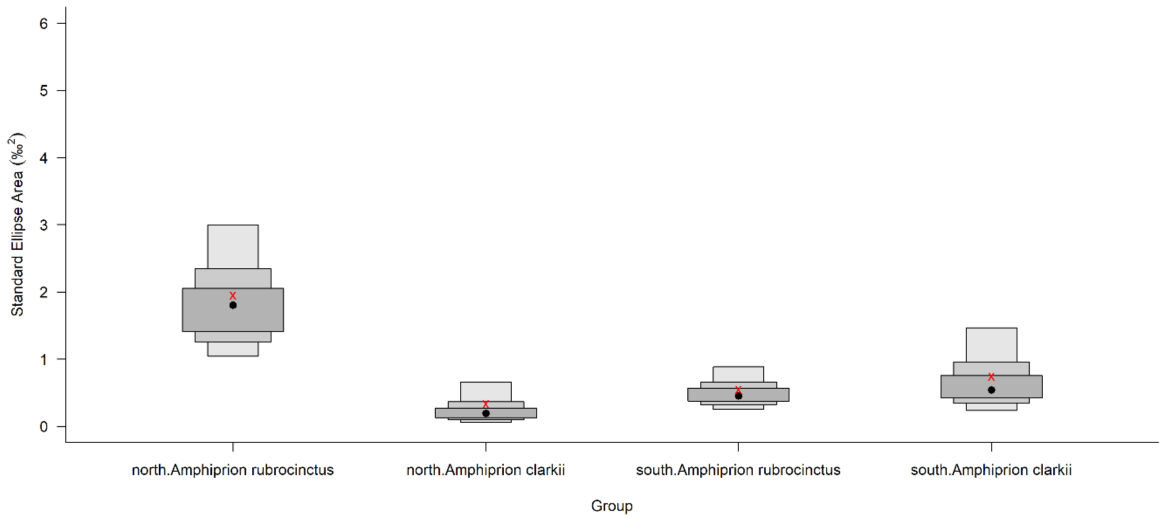


Supplementary Figure 22. Mean and standard error of standard ellipse areas for clownfish at different regions.

Supplementary Table 8. Isotopic niche overlap among different regions of clownfish species for standard ellipses (40%).

| | **Percent Niche Overlap (%)** | | | |
|---|---|---|---|---|
| | *A. clarkii - north* | *A. rubrocinctus - north* | *A. clarkii - south* | *A. rubrocinctus - south* |
| *A. clarkii - north* | | <0.01% | 0% | <0.01% |
| *A. rubrocinctus - north* | <0.01 | | 0% | 28% |
| *A. clarkii - south* | 0.00 | 0.00 | | 0% |
| *A. rubrocinctus - south* | <0.01 | 0.56 | 0.00 | |

**Ethanol preservation for stable isotopes**

Methods: Muscle from three adult and three juvenile individuals of each clownfish species were preserved in ethanol to determine whether ethanol preservation affected the stable isotope analysis. Ethanol treated samples were then placed in distilled water for 30min, oven dryed, and pulverized to a powder. Samples were placed in tin capsules and sent for stable isotope analysis as were other samples. Ethanol treated and untreated samples were compared using a paired t-test.
Results: There were no differences in ethanol-preserved versus nonpreserved clownfish samples for $\delta^{13}C$ values ($p = 0.54$), but ethanol samples had 0.34 higher $\delta^{15}N$ values than nonpreserved samples ($p < 0.01$). Therefore, ethanol affected $\delta^{15}N$ values only in clownfish.

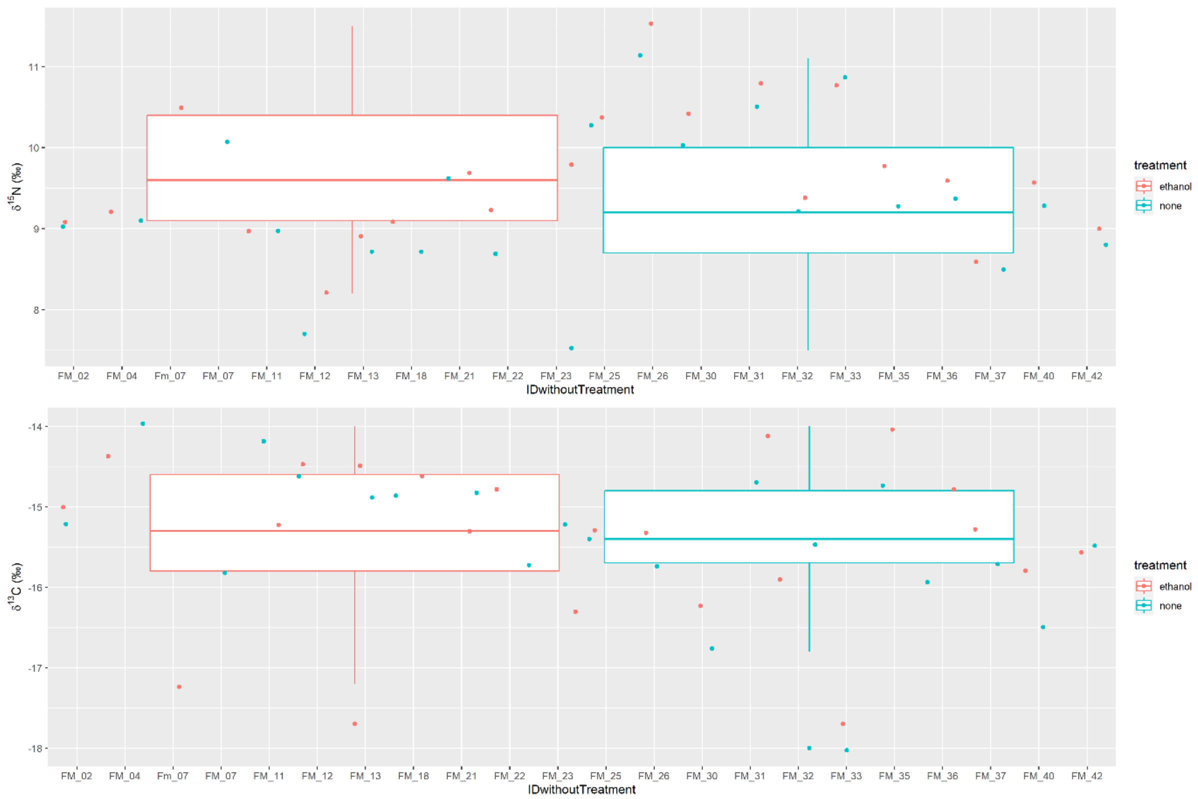


Supplementary Figure 23. Differences in isotopic values of carbon and nitrogen for ethanol-preserved and nonpreserved samples of clownfish.

#### *MICROBIOME DIVERSITY*

Supplementary Table 9. Output of all statistical analyses for microbiomes. PERMANOVA = permutational analysis of variance; LM = linear model.

| Response Variable | Model | Predictor variable | Factor Type | df | Model Statistic | Test-value | p-value |
|---|---|---|---|---|---|---|---|
| Microbiome of clownfish only | PERMANOVA | Species | fixed | 2 | F-value | 1.05 | 0.15 |
| (community) | | Life stage | fixed | 2 | F-value | 1.09 | 0.06 |
| | | Species x Life stage | interaction | 6 | F-value | 2.80 | 0.03* |
| | | | | | Sample size | n = 89 | |
| Microbiome of all samples | PERMANOVA | Species | fixed | 4 | F-value | 1.79 | <0.0001* |
| (community) | | Tissue Type | fixed | 3 | F-value | 2.10 | <0.0001* |
| | | Species x Tissue Type | interaction | 12 | F-value | 1.58 | <0.0001* |
| | | | | | Sample size | n = 136 | |
| Species Richness for Microbiome of clownfish only | LM | Species | fixed | 2 | F-value | 0.77 | 0.47 |
| log-transformed | | Life stage | fixed | 2 | F-value | 1.69 | 0.19 |
| | | Species x Life stage | interaction | 2 | F-value | 0.76 | 0.47 |
| | | R-squared = 0.073 | | | Sample size | n = 89 | |
| Shannon Diversity for Microbiome of clownfish only | LM | Species | fixed | 2 | F-value | 0.34 | 0.71 |
| untransformed | | Life stage | fixed | 2 | F-value | 0.75 | 0.48 |
| | | Species x Life stage | interaction | 2 | F-value | 0.60 | 0.55 |
| | | R-squared = 0.040 | | | Sample size | n = 89 | |
| Species Richness for all samples | LM | Species | fixed | 4 | F-value | 11.68 | <0.0001* |
| log-transformed | | Tissue Type | fixed | 3 | F-value | 19.23 | <0.0001* |
| | | Species x Tissue Type | interaction | 5 | F-value | 1.37 | 0.24 |
| | | R-squared = 0.475 | | | Sample size | n = 136 | |
| Shannon Diversity for all samples | LM | Species | fixed | 4 | F-value | 9.16 | <0.0001* |
| untransformed | | Tissue Type | fixed | 2 | F-value | 23.55 | <0.0001* |
| | | Species x Tissue Type | interaction | 2 | F-value | 2.71 | 0.023* |
| | | R-squared = 0.496 | | | Sample size | n = 136 | |

For shannon diversity indeces, there was no difference in life stage or species for fish only samples and there was no interaction (p > 0.19). Shannon diversity was different for species (p < 0.001) and tissue type (p < 0.001), and there was a significant interaction (p = 0.023). The mucus of *E. quadricolor* had the highest diversity and gills of *A. rubrocinctus* had the lowest diversity. There were no other differences in diversity although gills tended to have the lowest diversity overall.

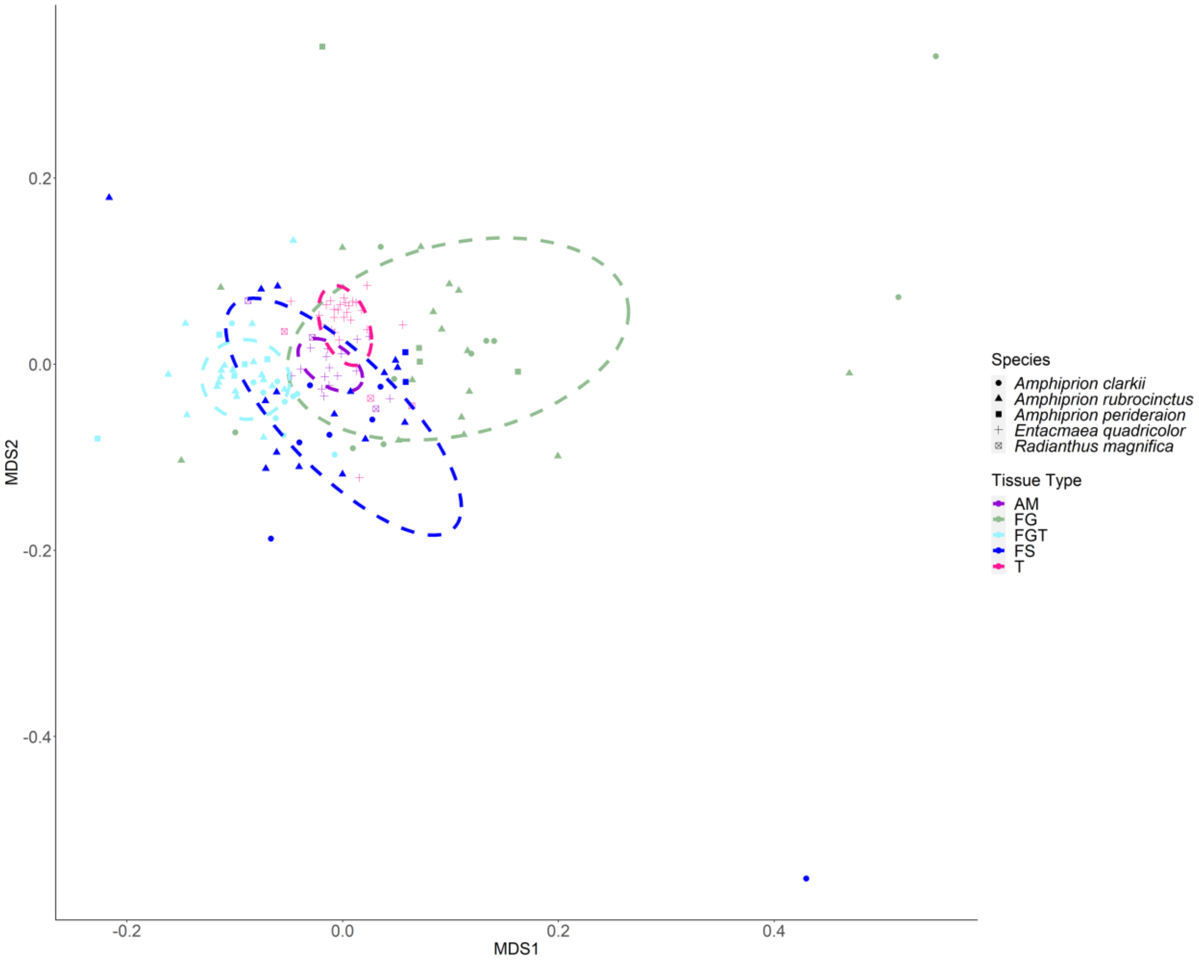


Supplementary Figure 24. The clusters of microbiome by tissue types (colors and ellipses) by species (shapes).

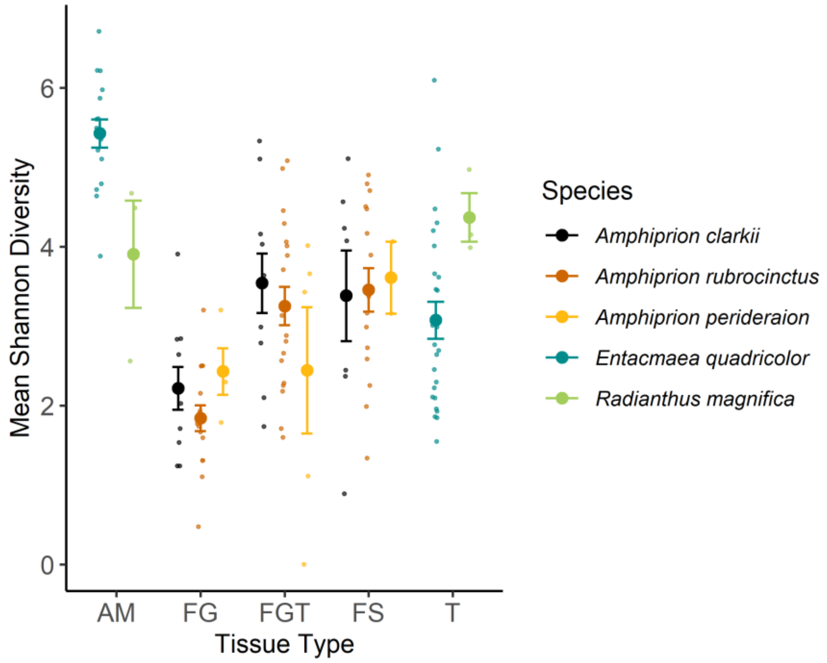


Supplementary Figure 25. Mean Shannon diversity of microbiome for all samples by species and tissue type.

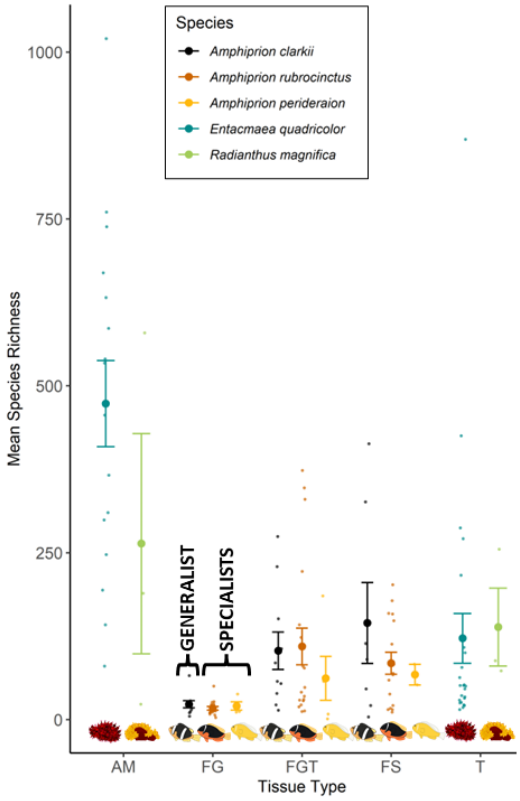


Supplementary Figure 26. Mean species richness of microbiome for all samples by species and tissue type.

Discussion:

Why prior hypotheses and directions of color pattern research on clownfish do not properly explain their ultimate function:

Prior to Gaboriau et al. (2025), other studies had been unable to recover a pattern linking clownfish coloration evolution to their host sea anemones (Merilaita & Kelley 2018, Salis et al. 2018, Salis et al. 2022, Hayashi et al. 2024). As a result, research that focused on explaining clownfish color pattern evolution and function turned towards hypotheses related to social function and species recognition. These broadly predicted that sympatric species should evolve sufficiently different color patterns in order to recognize conspecifics and discourage inter-specific associations. As an extension, intraspecific variation in color-pattern was hypothesized to communicate within group social status. A rich body of literature now highlights that clownfishes can discriminate among species using differences in the number of white vertical body bars, species with similar bar patterns occur less often in sympatry than expected by random chance, and that UV light reflectance may be important in communicating status within a social hierarchy. These are clear empirical color-pattern functions. Yet these functions are unable to broadly account for clownfish color-pattern evolution across the clade or explain why the host sea anemones are generating convergent clownfish phenotypes. We thus interpret these to be secondary, rather than ultimate, functions of clownfish color-patterns.

Only one previous study (Merilaita and Kelly 2018) proposed that clownfish color patterns serve protective functions. Using an older clownfish-sea anemone host-use matrix from Fautin and Allen (1992), they recovered correlations between vertical bar number, host tentacle length, host number, and anemone toxicity. Merilaita and Kelly (2018) hypothesized that ancestral clownfish used multiple white bars to effectively hide among anemone tentacles, and then later used vertical bars as an aposematic warning after evolving associations with highly toxic anemone species with short tentacles in the genus *Sticodactyla*. While we agree with Merilaita and Kelly in the hypothesis that clownfish color patterns are ultimately protective, our interpretation of color pattern function differs in key ways, and with the benefit of a fully revised host-use matrix and overwhelming signal that the hosts are responsible for color-pattern evolution (Gaboriau et al. 2025). In contrast to Merilaita and Kelly (2018), our predator vision modeling data show that vertical bars in clownfishes are highly visible to potential predators within their hosts, aligning with aposematic or disruptive coloration hypotheses for host generalists, but at odds with the notion that white bars can effectively be used as camouflage among host tentacles. Instead, we find the loss of bars and the evolution of base body colors that closely match their hosts tentacles leads to background matching phenotypes in host specialists.
